# Convergent anti-MRSA potency across compositionally distinct essential oils: a chemotype similarity index for strain-dependent chemistry–activity analysis

**DOI:** 10.64898/2026.07.02.736015

**Authors:** Atul Bhat, Angela Sherry

## Abstract

Antimicrobial resistance represents a continuing threat to clinical infection management, with methicillin-resistant *Staphylococcus aureus* (MRSA) and multidrug-resistant *Escherichia coli* identified by the World Health Organization as priority pathogens. This study evaluated the antimicrobial activity, synergistic potential, and chemical composition of six plant-derived preparations — three ethanolic extracts (nettle, thyme, rosemary) and three essential oils (lavender, lemongrass, doTERRA Peace blend) — against MRSA, methicillin-sensitive *S. aureus* (MSSA), and *E. coli* K-12 by disc diffusion, broth microdilution, post-exposure culturability, antimicrobial interactions assessed by checkerboard assay, and GC-MS profiling. Disc diffusion produced no interpretable zones of inhibition for any plant preparation tested, however broth microdilution revealed reproducible inhibitory activity within published ranges across the panel. Three essential oils achieved a median Minimum Inhibitory Concentration (MIC) of 0.39 mg/mL against MRSA despite presenting compositionally distinct chemotypes — lavender was linalool-dominated (61% combined), lemongrass was citral-dominated (76%), and the doTERRA blend was sesquiterpene-rich. Rosemary ethanolic extract achieved the same potency (0.39 mg/mL) against MSSA. No preparation produced a bactericidal reduction (≥3 log₁₀ CFU mL⁻¹) at any timepoint, with all reductions transient and recovering by 24 hours. Checkerboard combinations of plant preparations with vancomycin and ciprofloxacin were uniformly classified, according to the Fractional Inhibitory Concentration Index (FICI), as indifference/no interaction, attributable in part to inoculum-mediated effects on vancomycin MIC. To analyse the relationship between chemical composition and antimicrobial outcomes, we introduce a Chemotype Similarity Index (CSI), a chemometric framework quantifying pairwise compositional similarity between essential oils by Pearson correlation and relating it to log₂-MIC differences across strains. CSI revealed a strain-dependent chemistry–activity relationship — convergent against MRSA, monotonic against MSSA, and absent against E. coli — indicating that compositional similarity predicts antimicrobial outcomes on a strain-specific basis. The convergence of three chemotypically divergent essential oils with the same anti-MRSA potency suggested a shared membrane-disrupting mechanism operating through distinct chemical routes. Although exploratory at this scale, the CSI framework provides a reusable analytical scaffold for linking phytochemical composition to antimicrobial activity, and identifies the MRSA convergence as a specific direction for mechanistic investigation into the development of plant-derived antimicrobial adjuncts.

## 1. INTRODUCTION

Antimicrobial Resistance (AMR) is an escalating threat to global health that undermines the effectiveness of modern antimicrobial therapy. In 2019, AMR was directly responsible for approximately 1.27 million deaths worldwide, with a greater burden associated with resistant infections overall (Murray *et al*., 2022). This burden reflects both the prevalence of resistant pathogens and the cumulative effect of overlapping resistance mechanisms that reduce the efficacy of multiple antibiotic classes (Blair *et al*., 2015; Murray *et al.,* 2022). Among clinically significant pathogens, *Staphylococcus aureus* and *Escherichia coli* remain central contributors to AMR-associated morbidity and mortality. Methicillin-resistant *S. aureus* (MRSA) persists in hospital and community settings, driven by acquisition of the *mecA* gene encoding penicillin-binding protein PBP2a (Chambers and DeLeo, 2009; Lakhundi and Zhang, 2018). Recent structural evidence demonstrates that resistance in *S. aureus* is adaptable at the molecular level, reinforcing its clinical relevance (Alexander *et al*., 2023). Multidrug-resistant *E. coli* has emerged as a leading cause of difficult-to-treat infections through extended-spectrum β-lactamase production and plasmid-mediated resistance (Bush and Bradford, 2016). Resistance in Gram-negative bacteria is often compounded by reduced membrane permeability and efflux activity (Blair *et al*., 2015), and genomic surveillance confirms that resistant *E. coli* populations are diverse and transmissible (Brouwer *et al*., 2023; Maldonado *et al*., 2024). These mechanisms rarely act in isolation; their combined effects produce multidrug-resistant phenotypes that reduce treatment options and worsen clinical outcomes (Blair *et al*., 2015; Murray *et al*., 2022). Resistance to first-line and last-resort antibiotics has intensified the need for alternative or adjunct antimicrobial strategies (Tacconelli *et al*., 2018).

Medicinal plants are a significant source of biologically active compounds with antimicrobial potential, with recent reviews highlighting their relevance against pathogens prioritised by the World Health Organization (Iskandar *et al*., 2025). Phytochemicals such as phenolics and terpenoids act through disruption of membrane integrity and interfere with cellular processes (Cowan, 1999; Di Pasqua *et al*., 2007). Essential Oil (EO) components can induce structural damage to bacterial membranes, resulting in leakage of intracellular contents and loss of cellular viability (El-Sakhawy *et al*., 2026; Falleh, 2025). In this study, nettle, thyme and rosemary were selected based on their phytochemical composition and mechanistic relevance. Nettle contains phenolic compounds associated with antibacterial activity (Gülçin *et al*., 2004); thyme and rosemary are rich in terpenoids such as thymol and carvacrol, which disrupt bacterial membranes and increase permeability (Ultee *et al*., 1999; Marchese *et al*., 2016; Khwaza and Aderibigbe, 2025). Lavender EO and Lemongrass EO demonstrate antimicrobial potential through volatile compounds such as linalool and citral (Cavanagh and Wilkinson, 2005; Guo *et al*., 2021). Antimicrobial activity is highly dependent on concentration, formulation, and environmental conditions rather than being intrinsic to the compound itself (Bakkali *et al*., 2008; Van de Vel *et al*., 2019). This variability, combined with inconsistent methodologies, limits the comparability of findings across studies (Balouiri *et al*., 2016).

Extraction methodology determines the chemical composition and biological activity of plant-derived preparations. Ethanol is widely used because it extracts a broad range of phytochemicals (Dai and Mumper, 2010), but extraction conditions including solvent ratios and processing parameters can alter phytochemical yield and composition (Do *et al*., 2014). Differences in extraction protocols lead to inconsistent antimicrobial outcomes, making it difficult to compare findings across studies.

The potential for plant-derived compounds to enhance antibiotic efficacy is a key area of interest in AMR research. Resistance mechanisms such as efflux pump activity and reduced permeability limit antibiotic effectiveness (Blair *et al*., 2015), and compounds capable of disrupting these mechanisms may restore antimicrobial susceptibility. Carvacrol enhances antibiotic uptake in *E. coli* (Lv *et al*., 2011), and thymol improves β-lactam efficacy against resistant strains (Nascimento *et al*., 2000). Reviews of essential oil and antimicrobial combinations emphasise the heterogeneity of reported interaction outcomes (Soulaimani, 2025). Essential oils may act through multiple mechanisms simultaneously, including membrane disruption, efflux-mediated effects, interference with cell-to-cell signalling and biofilm-associated behaviour (Tang *et al*., 2020; Khwaza and Aderibigbe, 2025). These multi-target effects are relevant for multidrug-resistant pathogens, where single-target antibiotics often fail. Synergistic interactions vary with bacterial strain, compound concentration, and experimental conditions (Langeveld *et al*., 2014). The Fractional Inhibitory Concentration Index (FICI) provides a standardised method for evaluating interactions (Odds, 2003) but remains underutilised in many experimental designs.

Beyond interaction testing, the standard activity assays themselves carry interpretive limits. Minimum Inhibitory Concentration (MIC) assays are widely used to assess antimicrobial activity but provide limited mechanistic insight. MIC values indicate growth inhibition but do not distinguish between bacteriostatic and bactericidal effects (Andrews, 2001), nor do they capture the dynamics of bacterial killing. Reliance solely on endpoint MIC data limits ‘whole system’ interpretation.

Gas chromatography-mass spectrometry (GC-MS) is widely used to characterise essential oils and identify key constituents (Adams, 2007). Composition alone does not prove biological relevance. Prior work has attributed antimicrobial activity to specific dominant compounds without testing them in isolation against the target organism (Marchese *et al*., 2016). GC–MS profiling combined with whole-preparation assays can identify the dominant constituent classes within a preparation and the antimicrobial activity of the preparation as a whole, but cannot establish which compound or which combination, is responsible for the observed activity. This study presents compositional and antimicrobial data as complementary observations that generate mechanistic hypotheses for subsequent isolated-compound testing, rather than as evidence of direct causation between specific constituents and observed activity.

Research on plant-derived antimicrobials is constrained by methodological inconsistency, limited integration of phytochemical and microbiological data, and insufficient evaluation of synergistic interactions, even as recent reviews emphasise the urgency of accelerating EO research against priority resistant pathogens (Iskandar *et al*., 2025). Few studies combine controlled extraction methodologies with quantitative antimicrobial and synergy analyses across Gram-positive and Gram-negative pathogens. Given the global burden of AMR and the complexity of resistance mechanisms (Murray *et al*., 2022; Blair *et al*., 2015), addressing these limitations is essential for advancing plant-derived antimicrobials towards clinically meaningful application.

The aim of this study was to evaluate the antimicrobial activity, synergistic potential, and phytochemical composition of selected medicinal plant extracts and essential oils against clinically relevant bacterial pathogens, and to develop a quantitative framework relating essential-oil composition to antimicrobial outcome across strains.

This aim was addressed in two parts. First, the inhibitory activity of the ethanolic plant extracts and essential oils was determined against *Staphylococcus aureus*, MRSA, and *Escherichia coli*, and their combinations with conventional antibiotics were characterised for interactions classifiable by Fractional Inhibitory Concentration Index (FICI) thresholds.

Second, quantitative compositional similarity between the essential oils, derived from GC-MS profiles, was related to their relative antimicrobial activity to test whether the strength and direction of this relationship varied by bacterial strain.

## 2. MATERIALS AND METHODS

### 2.1 Plant Material and Essential Oils

Dried plant materials were obtained from commercial suppliers: organic rosemary (*Rosmarinus officinalis* L.), Hatton Hill Organic, nettle leaf (*Urtica dioica* L.), Manor Springs, and organic thyme (*Thymus vulgaris* L.), Spice Planet Ltd. Essential oils included lavender (*Lavandula angustifolia*), Tisserand Aromatherapy, lemongrass (*Cymbopogon flexuosus*), Naissance, and doTERRA Peace blend, doTERRA International, used as supplied. Dried materials were ground (CHP61 Kenwood Ltd, UK) and sieved through a 250 µm mesh.

### 2.2 Preparation of Ethanolic Extracts from Plant Materials

Extracts were prepared by maceration of 17 g dried material in 170 mL of 95% ethanol (1:10 w/v) at 100 rpm for 96 hours at room temperature. Extracts were filtered through Whatman No.1 filter paper, then concentrated under reduced pressure (∼25 mbar) at 40°C by rotary evaporation (Sticher, 2008) and reconstituted in DMSO (Sigma-Aldrich, Gillingham, UK) at 100 mg/mL.

### 2.3 Preparation of Essential Oil Solutions

Essential oil stocks were prepared at 100 mg/mL in DMSO and diluted to working concentrations in brain heart infusion broth (BHI, Neogen, Ayr, Scotland) for all assays.

### 2.4 Bacterial Strains and Culture Conditions

Three strains were used: *Staphylococcus aureus* (DSM 20231), methicillin-resistant *S. aureus* MRSA (laboratory clinical isolate, not formally characterised; findings preliminary), and *Escherichia coli* K-12 (DSM 3925). The MRSA isolate has been maintained by extended laboratory sub-culture and may have undergone phenotypic drift; results require validation against a characterised reference strain. Strains were cultured in BHI broth at 37°C for 18-24 hours and adjusted to a 0.5 McFarland standard for all assays. All procedures were conducted aseptically.

### 2.5 Disc Diffusion Assay

Antimicrobial activity was screened by disc diffusion following the Clinical and Laboratory Standards Institute (CLSI) guidelines (2022). Sterile filter paper discs impregnated with 10 µL of each preparation were placed on BHI agar plates inoculated with 200 µL of each strain adjusted to a 0.5 McFarland standard and incubated at 37°C for 24 hours. Zones of inhibition were measured in millimetres. Vancomycin and ciprofloxacin served as positive controls for Gram-positive and Gram-negative strains; DMSO was the negative control. Assays were conducted in triplicate. Specific concentrations and preparations are listed in Supplementary material 1 (SM1) [Table 1].

**Table 1.** Compound identification via GC-MS in three essential oil preparations.

| No. | Compound | RT (min) | HIT | Rel. | LAV (%) | LEM (%) | dT (%) |
| --- | --- | --- | --- | --- | --- | --- | --- |
| 1 | α-Pinene | 6.23 | 931 | **R** | 0.34 | — | 6.46 |
| 2 | Camphene | 6.52 | 882 | **P** | 0.55 | 2.33 | — |
| 3 | Eucalyptol (1,8-cineole) | 7.86 | 762 | **T** | 1.99 | 0.8 | 0.3 |
| 4 | Linalool | 8.89 | 949 | **R** | 24.8 | — | 7.3 |
| 5 | cis-p-Menth-8-en-1-ol | 8.97 | 938 | **R** | 2.7 | — | 2.75 |
| 6 | endo-Borneol | 10.09 | 935 | **R** | 1.13 | — | — |
| 7 | Terpinen-4-ol | 10.23 | 861 | **P** | 7.61 | 0.91 | 1.79 |
| 8 | (-)-cis-Isopiperitenol | 10.77 | 901 | **R** | — | 1.41 | — |
| 9 | Neral (citral, Z-isomer) | 11.02 | 894 | **P** | — | 32.58 | — |
| 10 | Linalyl acetate | 11.12 | 954 | **R** | 36.13 | — | 14.14 |
| 11 | Geraniol | 11.17 | 857 | **P** | — | 1.68 | — |
| 12 | Neral (secondary aldehyde) | 11.31 | 832 | **P** | — | 1.53 | — |
| 13 | Geranial (E-2,6-octadienal) | 11.44 | 944 | **R** | — | 43.33 | — |
| 14 | Lavandulyl acetate | 11.59 | 954 | **R** | 5.81 | — | — |
| 15 | Copaene | 13.01 | 871 | **P** | — | 1.64 | 0.42 |
| 16 | cis-α-Bisabolene | 13.55 | 823 | **P** | 0.95 | — | — |
| 17 | β-Caryophyllene | 13.62 | 824 | **P** | 7.9 | 3.9 | 5.28 |
| 18 | β-Longipinene | 14.29 | 792 | **T** | 0.33 | — | 3.03 |
| 19 | Germacrene D | 14.4 | 865 | **P** | 0.75 | — | 2.95 |
| 20 | Caryophyllene oxide | 15.68 | 818 | **P** | 1.03 | 1.76 | 1.06 |
| 21 | Cadinene-type sesquiterpene | 17.29 | 875 | **P** | — | — | 2.82 |
| 22 | Aromadendrene-type sesquiterpene | 17.57 | 846 | **P** | — | — | 5.86 |
| 23 | (E)-Nerolidol | 17.97 | 939 | **R** | — | — | 8.44 |

{Reliability key: R = reliable identification (NIST 2017 HIT score $\geq 900$ ); P = probable (HIT 800–899); T = tentative (HIT 700–799). Compounds with HIT < 700 were excluded from analysis. LAV = Lavender essential oil; LEM = Lemongrass essential oil; dT = doTERRA Peace blend essential oil; RT = Retention Time (in minutes). The dash (—) indicated compound not detected in that preparation. Citral comprised two geometric isomers, neral (Z) and geranial (E), which elute as separate peaks; peak 9 is the neral isomer and peak 13 is geranial (reported by the library as 2,6-octadienal, 3,7-dimethyl-, (E)-). The combined citral content cited in the text ( $\approx 76\%$ ) is the sum of peaks 9 and 13 (32.6% + 43.3%); the 32.6% value refers to peak 9 (neral) alone. Peak 12 (1.5%; HIT 832) is a minor, separately eluting aldehyde returned by the library as neral; given the low match score and the dominant neral assignment to peak 9, it was reported as a minor secondary aldehyde and was not included in the citral total.}

### 2.6 Minimum Inhibitory Concentration (MIC) determination

MIC were determined by broth microdilution in 96-well plates (Wiegand et al., 2008). Each plate contained six plant preparations tested in two-fold serial dilution across columns 1–8 (final concentrations 50–0.39 mg/mL), with shared control columns for blank, solvent, and growth references. Bacterial inocula were prepared to a 0.5 McFarland standard (verified at optical density at 600 nm [OD₆₀₀] 0.08–0.10 against a BHI blank), then diluted 1:200 in broth; 10 µL was added per test well to give ∼5 × 10⁵ CFU/mL final density (total test volume 110 µL). OD₆₀₀ was recorded at 0 h and 24 h after 37 °C incubation on a Tecan Infinite 200 Pro reader (Tecan, Switzerland). Relative growth (RG) was calculated from the change in optical density over incubation (ΔOD = OD₆₀₀ at 24 h − OD₆₀₀ at 0 h) as (ΔOD sample − ΔOD blank) ÷ (ΔOD growth control − ΔOD blank) × 100; MIC was the lowest concentration at which RG ≤ 10%. Full plate layout and control column assignments are provided in SM2. Results were reported as medians across three biological replicates (Andrews, 2001).

### 2.7 Post-exposure Culturability Assay

Lavender essential oil, rosemary ethanolic extract, and thyme ethanolic extract were selected based on their MIC profiles against MRSA, MSSA, and *E. coli,* respectively. Inoculum was prepared by adjusting overnight BHI cultures to a 0.5 McFarland standard, verified by absorbance at 600 nm against a BHI blank (OD₆₀₀ 0.08–0.10). Standardised suspensions were diluted 1:10 into the assay, yielding ∼1 × 10⁷ CFU mL⁻¹ — approximately 20-fold higher than the MIC inoculum (Section 2.6), consistent with time-kill convention. Preparations were combined with inoculum at 1× and 2× MIC (extracts). At 0, 2, 6, and 24 hours, aliquots were diluted in phosphate-buffered saline (PBS) and plated by drop-plate (10 µL drops) on BHI agar across serial ten-fold dilutions; colonies were counted after 24 hours at 37°C and expressed as log₁₀ CFU mL⁻¹. CFU values are the mean of valid (non-TMTC [too many to count], non-zero) replicates at the optimal countable dilution. Without a validated neutralisation step, data are presented as exploratory post-exposure culturability measures rather than time-kill kinetics. Bactericidal activity was defined as ≥3 log₁₀ CFU mL⁻¹ reduction from baseline (Wiegand *et al*., 2008).

### 2.8 Checkerboard Fractional Inhibitory Concentration Index (FICI) Synergy Assay

Four combinations were assessed in triplicate: 1) lavender with vancomycin against MRSA; 2) lemongrass with vancomycin against MRSA; 3) rosemary with vancomycin against MSSA; and 4) lavender with ciprofloxacin against *E. coli* K-12. Antibiotic working stocks were prepared at 2.2× the target final concentration to account for in-well dilution (50 µL antibiotic in a 110 µL final well): vancomycin 132 µg/mL and ciprofloxacin 1.10 µg/mL, giving final column-1 concentrations of 60 µg/mL and 0.50 µg/mL respectively (each 2× the standalone MIC of 30 µg/mL and 0.25 µg/mL). Extract starting concentrations were 0.78 mg/mL (2× MIC) for combinations 1–3 and 1.56 mg/mL (1× MIC) for combination 4, limited by the 1% DMSO threshold. Plates were incubated at 37°C for 24 hours; OD₆₀₀ and RG calculated as in Section 2.6. Fractional Inhibitory Concentration Index (FICI) = (MIC extract,combination ÷ MIC extract,alone) + (MIC antibiotic,combination ÷ MIC antibiotic,alone), classified according to Pillai *et al*. (2005) and Odds (2003): ≤ 0.5 synergy; 0.5–1.0 additive; 1.0–4.0 indifference; > 4.0 antagonism.

### 2.9 Qualitative Phytochemical Screening

The ethanolic plant extracts were screened in triplicate for flavonoids (Shinoda test; Harborne, 1998), alkaloids (Wagner’s reagent; Sreevidya and Mehrotra, 2003), phenolics (ferric chloride test; Dai and Mumper, 2010), and saponins (froth test; Sofowora, 1993). Reagent compositions, volume ratios, and per-replicate observations are provided in SM3.

### 2.10 Gas Chromatography–Mass Spectrometry (GC–MS)

All six preparations were profiled at 1 mg/mL in DMSO on a Thermo Scientific Trace 1300 GC coupled to an ISQ 7000 single quadrupole mass spectrometer with TriPlus RSH autosampler (Thermo Fisher Scientific, USA). Separation used a TG5-MS capillary column (30 m × 0.25 mm × 0.25 µm) with helium at 1 mL/min. Temperature programme: 40°C for 1 minute, 10°C/min ramp to 300°C, 2-minute hold; injector 280°C; 1 µL injection; split 15:1; EI ionisation at 50–650 m/z. Compounds were identified against the NIST 2017 spectral library; match scores below 700 were excluded, 700–799 reported as tentative (T), 800–899 probable (P), and ≥ 900 reliable (R). Peak areas were normalised to the total integrated peak area and expressed as a percentage of total ion current for each preparation.

### 2.11 Chemotype Similarity Index (CSI)

#### 2.11.1 Compositional vector construction

For each essential oil profiled by GC-MS (Section 2.10), the relative percentage abundance of every compound identified above the NIST 2017 match score threshold of 700 was extracted from the total ion chromatogram and normalised to the total identified compound area, yielding a fractional compositional vector for each preparation. The union of compounds appearing within the top ten constituents of any of the three essential oils was constructed, yielding 23 unique compounds (Table 1) and provided in SM4. Compositional vectors for each preparation were aligned across this 23-compound space, with compounds absent from a given preparation assigned a value of zero.

#### 2.11.2 Pairwise compositional similarity

Chemical similarity between each pair of essential oils was quantified using Pearson product-moment correlation of their compositional vectors across the unified compound space, constructed from GC-MS relative abundances as reported, with compounds absent from a given preparation encoded as zero. No additional per-vector renormalisation was applied. The Pearson coefficient (r) ranges from −1 (compositionally opposite) through 0 (compositionally unrelated) to +1 (compositionally identical), and reflects whether high-abundance compounds in one preparation are also high-abundance in another. Pearson correlation was selected over alternative similarity metrics (Bray-Curtis, cosine similarity, Spearman rank) as it preserves both the direction and magnitude of compositional correspondence and is the metric most commonly applied in GC-MS chemometric comparisons (Adams, 2007).

A summary statistic, mean CSI, was calculated as the arithmetic mean of the three pairwise Pearson correlations across the three essential oil preparations. Mean CSI provides a panel-level descriptor of overall chemical heterogeneity within the essential oil collection: values approaching +1 indicate compositional convergence; values approaching 0 indicate compositional independence; values approaching −1 indicate compositional divergence.

#### 2.11.3 Chemistry–activity linkage analysis

For each bacterial strain, the relationship between chemical similarity and antimicrobial similarity was examined by plotting pairwise compositional correlations against the absolute log₂-MIC difference between the same pairs of preparations, derived from the median MIC values reported in Section 3.3. Log₂ transformation was applied because MIC values were generated by two-fold serial dilution; log₂ differences therefore correspond directly to the dilution steps separating preparations. A monotonic relationship — small log₂-MIC differences associated with high *r*, and vice versa — would indicate that compositional dominance predicts antimicrobial outcome. The absence of such a relationship would indicate that activity is not determined by which compounds are dominant.

#### 2.11.4 Implementation

Pearson correlations and log₂-MIC differences were calculated in Python 3.11 using NumPy 1.26. The CSI calculation script and worked calculation example are provided in SM4. CSI is presented as an exploratory descriptive framework based on n = 3 essential oil preparations; statistical inference on the correlation coefficients was not performed because the sample size precludes adequately powered hypothesis testing.

#### 2.11.5 Scope

CSI was applied only to the three essential oil preparations (lavender, lemongrass, doTERRA Peace blend). The three ethanolic extracts (nettle, thyme, rosemary) produced GC-MS profiles dominated by matrix compounds without identifiable botanical terpene constituents (Sections 2.10 and 3.6) and were therefore excluded from CSI analysis.

### 2.12 Statistical Analysis

All assays were conducted in triplicate (n = 3 replicates per preparation–strain combination). Post-exposure data are expressed as log₁₀ CFU mL⁻¹ relative to growth control baselines. Phytochemical screening results are qualitative.

## 3. RESULTS

### 3.1 Overview of the experimental dataset

Six plant-derived preparations were evaluated against MRSA, MSSA, and *E. coli* K-12 across six experimental approaches. The preparations comprised ethanolic extracts of nettle, thyme, and rosemary, and essential oils of lavender, lemongrass, and doTERRA Peace blend. Disc diffusion produced no interpretable zones of inhibition for any preparation against any strain, and broth microdilution was therefore adopted as the primary quantitative method (Sections 3.2 and 3.3). Post-exposure culturability was assessed for selected preparations against each strain (Section 3.4). Checkerboard synergy assays were conducted for four preparation–antibiotic combinations (Section 3.5). Chemical characterisation by phytochemical screening and GC-MS profiling is reported in Section 3.6.

### 3.2 Antimicrobial screening by disc diffusion

Antibiotic positive controls confirmed assay validity: vancomycin produced mean zones of 4.7 ± 0.6 mm (MRSA) and 4.3 ± 0.6 mm (MSSA), and ciprofloxacin produced 12.5 ± 0.9 mm against *E. coli* K-12 (Table 2). DMSO 1% (v/v) negative controls produced 0.0–1.7 mm readings attributable to mechanical disturbance during disc placement, not inhibition.

**Table 2.** Zone of inhibition (mm) produced by plant preparations and controls in disc diffusion assays against MRSA, MSSA, and *E. coli*.

| Preparation | Conc. | MRSA<br>Mean ± SD<br>(mm) | Range<br>(mm) | MSSA<br>Mean ± SD<br>(mm) | Range<br>(mm) | *E. coli*<br>K-12<br>Mean ± SD<br>(mm) | Range<br>(mm) |
| --- | --- | --- | --- | --- | --- | --- | --- |
| Vancomycin | 30 μg | 4.7 ± 0.6 | 4–5 | 4.3 ± 0.6 | 4–5 | NT | — |
| Ciprofloxacin | 5 μg | NT | — | NT | — | 12.5 ± 0.9 | 12–13.5 |
| DMSO<br>(negative) | 1% v/v | 1.7 ± 2.1 | 0–4 | 0.3 ± 0.6 | 0–1 | 0.0 ± 0.0 | 0 |
| Nettle EE | 50 mg/mL | 1.3 ± 2.3 | 0–4 | 0.3 ± 0.6 | 0–1 | 0.7 ± 1.2 | 0–2 |
| Nettle EE | 25 mg/mL | 0.0 ± 0.0 | 0 | NT | — | NT | — |
| Thyme EE | 50 mg/mL | NT | — | 0.0 ± 0.0 | 0 | NT | — |
| Thyme EE | 25 mg/mL | NT | — | 0.3 ± 0.6 | 0–1 | NT | — |
| Rosemary EE | 50 mg/mL | NT | — | 2.0 ± 2.0 | 0–4 | 5.7 ± 1.5 | 4–7 |
| Lavender EO | 10% w/v | 1.7 ± 1.5 | 0–3 | NT | — | 2.0 ± 3.5 | 0–6 |
| Lemongrass EO | 10% w/v | 0.0 ± 0.0 | 0 | 0.0 ± 0.0 | 0 | 0.0 ± 0.0 | 0 |
| doTERRA Peace<br>EO | 10% w/v | 5.3 ± 2.3 | 4–8 | NT | — | 1.3 ± 2.3 | 0–4 |

No plant preparation produced an interpretable zone; most measurements were indistinguishable from the 6 mm disc. Lemongrass produced 0.0 ± 0.0 mm against all three strains despite the lowest median MIC values by broth microdilution — 0.39 mg/mL against both MRSA and MSSA (Section 3.3, Table 3). The doTERRA Peace blend produced the largest mean zone of inhibition against MRSA at 5.3 ± 2.3 mm, with inter-replicate variability of 4–8 mm precluding reliable interpretation. Rosemary against *E. coli* produced the most reproducible phytochemical result at 5.7 ± 1.5 mm. Per-replicate measurements are provided in SM1 (Table 1).

**Table 3.** Minimum inhibitory concentrations of plant extracts and essential oils against clinically relevant bacterial pathogens.

| Sample | Type | MRSA<br>Median (range)<br>mg/mL | MSSA<br>Median (range)<br>mg/mL | *E. coli* K-12<br>Median (range)<br>mg/mL |
| --- | --- | --- | --- | --- |
| Nettle | EE | 1.56<br>(0.39–6.25) | 1.56<br>(0.39–6.25) | 3.13<br>(0.39–6.25) |
| Thyme | EE | 3.13<br>(0.78–3.13) | 3.13<br>(0.39–12.5) | 3.13<br>(0.39–3.13) |
| Rosemary | EE | 1.56<br>(0.39–12.5) | 0.39<br>(0.39–3.13) | 3.13<br>(0.39–3.13) |
| Lavender | EO | 0.39<br>(0.39–1.56) | 1.56<br>(1.56–6.25) | 1.56<br>(1.56–3.13) |
| Lemongrass | EO | 0.39<br>(0.39) | 0.39<br>(0.39–50) § | 0.78<br>(0.78–50) § |
| doTERRA Peace<br>blend | EO | 0.39<br>(0.39–6.25) | 3.13<br>(0.39–50) § | 12.5<br>(0.39–12.5) |

EE = ethanolic extract; EO = essential oil.

### 3.3 Minimum inhibitory concentration (MIC) results

MIC values for all six preparations against three strains are presented in Table 3; per-replicate data and worked relative growth calculations are provided in SM2 (Tables 1–4). All preparations demonstrated inhibitory activity within the tested range (0.39–50 mg/mL) against at least one of the strains.

**Table 4.** Overview of checkerboard assay combinations.

| Combination | Preparation | Type | Antibiotic | Strain | Highest extract | Highest antibiotic | n |
| --- | --- | --- | --- | --- | --- | --- | --- |
| 1 | Lavender | EO | Vancomycin | MRSA | 0.78 mg/mL (2× MIC) | 60.0 µg/mL (2× MIC) | 3 |
| 2 | Lemongrass | EO | Vancomycin | MRSA | 0.78 mg/mL (2× MIC) | 60.0 µg/mL (2× MIC) | 3 |
| 3 | Rosemary | EE | Vancomycin | MSSA | 0.78 mg/mL (2× MIC) | 60.0 µg/mL (2× MIC) | 3 |
| 4 | Lavender | EO | Ciprofloxacin | *E. coli* K-12 | 1.56 mg/mL (1× MIC)* | 0.50 µg/mL (2× MIC) | 3 |

EE = Ethanolic extract; EO = Essential Oil

The essential oils returned the lowest median MIC values across the panel. Lemongrass exhibited the most reproducible activity against MRSA, with identical MIC values in all three replicates (median 0.39 mg/mL). Lavender demonstrated broad-spectrum activity, returning median MIC values of 0.39 mg/mL against MRSA, 1.56 mg/mL against MSSA, and 1.56 mg/mL against *E. coli* — the lowest value recorded for any preparation tested against *E. coli*. The doTERRA Peace blend returned a comparably low median MIC against MRSA (0.39 mg/mL) yet reduced activity against *E. coli* (median 12.5 mg/mL).

Among the ethanolic extracts, rosemary returned the lowest median MIC against MSSA (0.39 mg/mL; range 0.39–3.13 mg/mL). Nettle showed moderate activity across all three strains (1.56 mg/mL against MRSA and MSSA; 3.13 mg/mL against *E. coli*). Thyme returned 3.13 mg/mL against all three microorganisms. All three ethanolic extracts retained inhibitory activity against *E. coli*. Vancomycin and ciprofloxacin controls produced fully concordant inhibition (Table 3; per-replicate values in SM6).

Inter-replicate variability was low for lavender across all three strains (CV <15%) and for lemongrass against MRSA (CV = 0%). Anomalous replicates yielding no inhibition at concentrations up to 50 mg/mL were recorded for lemongrass against MSSA and *E. coli,* and for the doTERRA blend against MSSA (SM2, Table 4). Median values are therefore considered a reliable summary statistic. Taken together, the MIC data demonstrate a consistent pattern in which essential oils outperformed ethanolic extracts against Gram-positive organisms, with the three essential oils each achieving 0.39 mg/mL against MRSA. This differential is explored in Section 3.6 and discussed in Sections 4.2 and 4.3.

### 3.4 Exploratory post-exposure culturability

Viable colony counts at 1× and 2× MIC over 24 hours are presented in Figure 1; calculated log₁₀ CFU mL⁻¹ values and the drop-plate conversion methodology are provided in SM5 (Tables 1-3). No preparation produced a ≥3 log₁₀ CFU mL⁻¹ reduction at any timepoint, the conventional threshold for bactericidal activity (Wiegand *et al*., 2008). All reductions were transient, with counts recovering to near-baseline by 24 hours.

**Figure 1.**
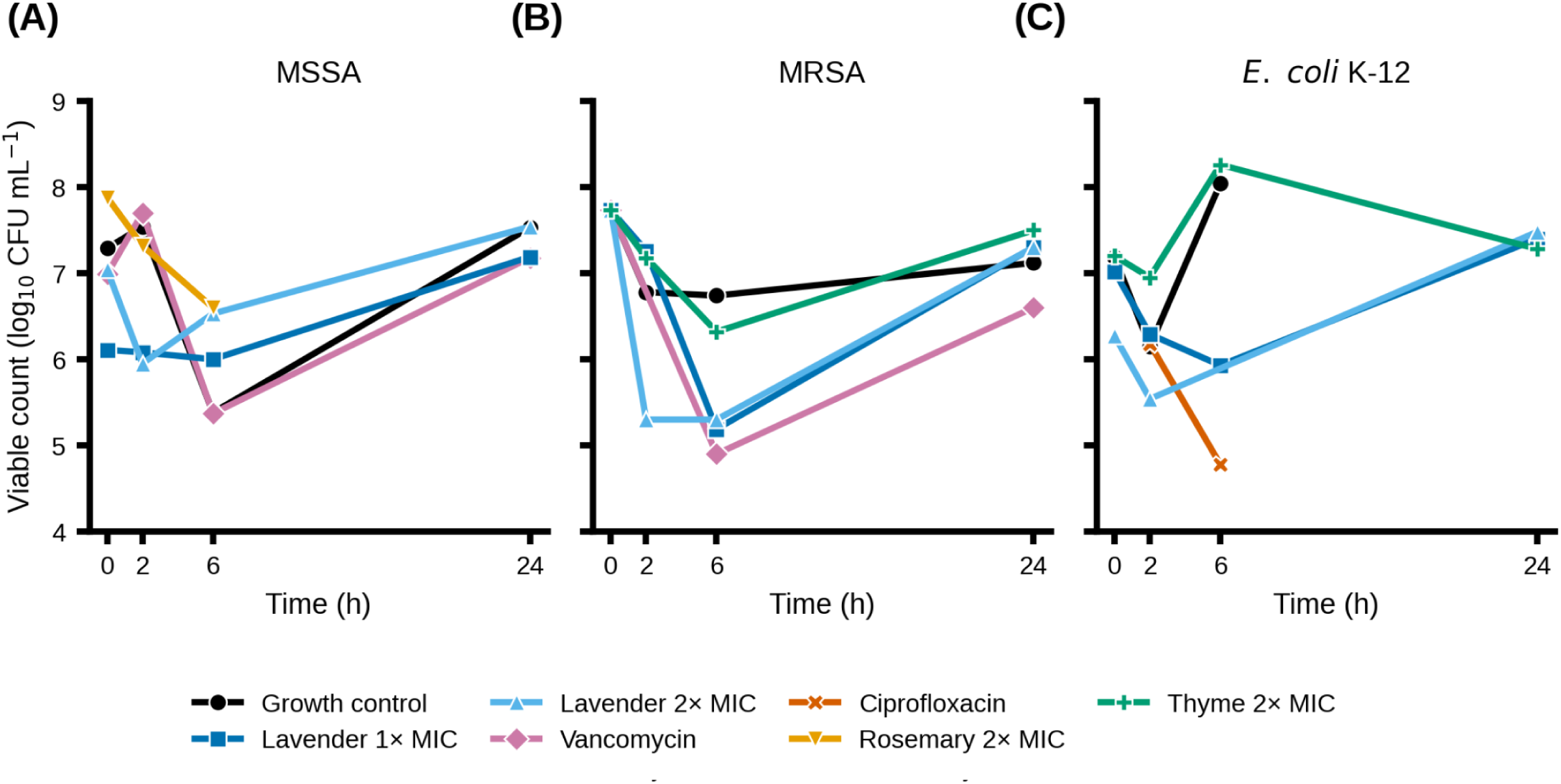
Post-exposure culturability of MSSA, MRSA, and *E. coli* K-12 following treatment with selected plant-derived preparations. Viable colony counts (log₁₀ CFU mL⁻¹) determined at 0, 2, 6, and 24 h post-exposure (n = 3 drop replicates per timepoint per condition). (A) MSSA; (B) MRSA; (C) *E. coli* K-12. Plant preparations: lavender essential oil at 1× and 2× MIC; rosemary ethanolic extract at 2× MIC (panel A only); thyme ethanolic extract at 2× MIC (panels B and C only). Antibiotic controls at standalone MIC: vancomycin (30 µg mL⁻¹) for MSSA and MRSA; ciprofloxacin (0.25 µg mL⁻¹) for *E. coli* K-12. Growth control: bacterial inoculum in BHI without treatment. The MRSA growth control 0 h value (7.73 log₁₀ CFU mL⁻¹) is used as the reference baseline in panel B because treatment wells were above the countable range at 0 h. Bactericidal activity was defined as a ≥3 log₁₀ CFU mL⁻¹ reduction relative to the 0 h baseline (Wiegand et al., 2008); no condition met this criterion at any timepoint. Growth-control variation between adjacent timepoints (e.g. MSSA at 6 h) reflects drop-plate sampling variation and lies within the technical noise of the assay. Excluded data points: MRSA vancomycin 2 h (anomalous, 8.95 log₁₀ CFU mL⁻¹); *E. coli* ciprofloxacin 0 h and 24 h (below detection); *E. coli* lavender 2× MIC 6 h, MSSA rosemary 2× MIC 24 h, and *E. coli* growth control 24 h (too many to count). Per-timepoint values and conversion methodology in SM5.

The most pronounced reductions were observed for lavender against MRSA (Figure 1B). Treatment wells for MRSA were above the countable range at 0h; the growth control 0h value of 7.73 log₁₀ CFU mL⁻¹ was used as the reference baseline. At 2× MIC, counts declined to 5.30 log₁₀ CFU mL⁻¹ by 2 hours (2.43-log reduction). At 1× MIC, a 2.54-log reduction was observed by 6 hours. In both cases, counts recovered to 7.30 log₁₀ CFU mL⁻¹ by 24 hours.

Against MSSA (Figure 1A), lavender at both 1× and 2× MIC produced reductions of approximately 1.2–1.3 log₁₀ CFU mL⁻¹ by 2–6 hours (relative to growth control 0h of 7.29), with full recovery by 24 hours. Rosemary at 2× MIC produced a 0.69-log reduction by 6 hours against MSSA; the 24-hour rosemary count was above the countable range at all dilutions and was excluded from Figure 1.

Against MRSA, thyme at 2× MIC produced a 1.41-log reduction at 6 hours (7.73 to 6.32 log₁₀ CFU mL⁻¹) with near-complete regrowth by 24 hours. Against *E. coli* (Figure 1C), thyme produced no measurable reduction. Lavender against *E. coli* at 2× MIC produced a 1.65-log reduction at 2 hours (GC 0h baseline 7.19 to 5.54); at 1× MIC, reductions reached 1.26 log₁₀ CFU mL⁻¹ at 6 hours, with full recovery by 24 hours.

Vancomycin maintained suppression of MRSA and MSSA at 6 and 24 hours, and ciprofloxacin produced complete *E. coli* suppression at 0 and 24 hours (below the drop-plate detection limit of ∼2 log₁₀ CFU mL⁻¹). An anomalous 8.95 log₁₀ CFU mL⁻¹ in the MRSA vancomycin control at 2 hours, and a single unresolvable 6h value for lavender 2× MIC against *E. coli*, were excluded as technical artefacts.

### 3.5 Checkerboard analysis and FICI classification

Checkerboard assays evaluated interactions across four two-compound combinations assessed across 12 plates (Table 4).

Growth controls were valid across all 12 plates (growth-control ΔOD 0.91–1.26; SM7, Table 2). Formal FICI values were not calculable in any combination, as inhibition boundaries could not be resolved in either reference control (Table 5); all four interactions were classified as ‘indifference’.

**Table 5.** FICI classification summary across all four combinations.

| Combination | Preparation | Antibiotic | Strain | GC valid | AB control | Extract MIC reached | FICI | Classification |
| --- | --- | --- | --- | --- | --- | --- | --- | --- |
| 1 | Lavender EO | Vancomycin | MRSA | Yes (3/3) | Not inhibited | No (93–98% at 2× MIC) | N/C | Indifference |
| 2 | Lemongrass EO | Vancomycin | MRSA | Yes (3/3) | Not inhibited | No (growth at 2× MIC) | N/C | Indifference |
| 3 | Rosemary EE | Vancomycin | MSSA | Yes (3/3) | Not inhibited | No (growth at 2× MIC) | N/C | Indifference |
| 4 | Lavender EO | Ciprofloxacin | *E. coli* | Yes (3/3) | Inhibited reps 1,3; partial rep 2 | No (82–105% at 1× MIC) | N/C | Indifference |

N/C = Non-calculable; Classification as per Odds (2003)

For combinations 1 (lavender + vancomycin vs MRSA), 2 (lemongrass + vancomycin vs MRSA), and 3 (rosemary + vancomycin vs MSSA), the vancomycin-alone reference row failed to show concentration-dependent inhibition across all nine replicates. At 60 µg/mL (2× MIC), relative growth ranged from 42–96%, indicating vancomycin did not inhibit growth at twice its standalone MIC (Column 1, SM7 Table 3). The extract-alone columns similarly showed no inhibition: lavender at 0.78 mg/mL returned 93–98% RG; lemongrass at 0.78 mg/mL returned 95–104%; rosemary at 0.78 mg/mL returned 82–99%. No combination matrix well demonstrated inhibition for combinations 1, 2 or 3. In combination 1, apparent inhibition at row A columns 1–2 in replicate 1 was an artefact of blank well contamination and was excluded (Figure 2; plate validity checks and exclusions in SM7).

**Figure 2.**
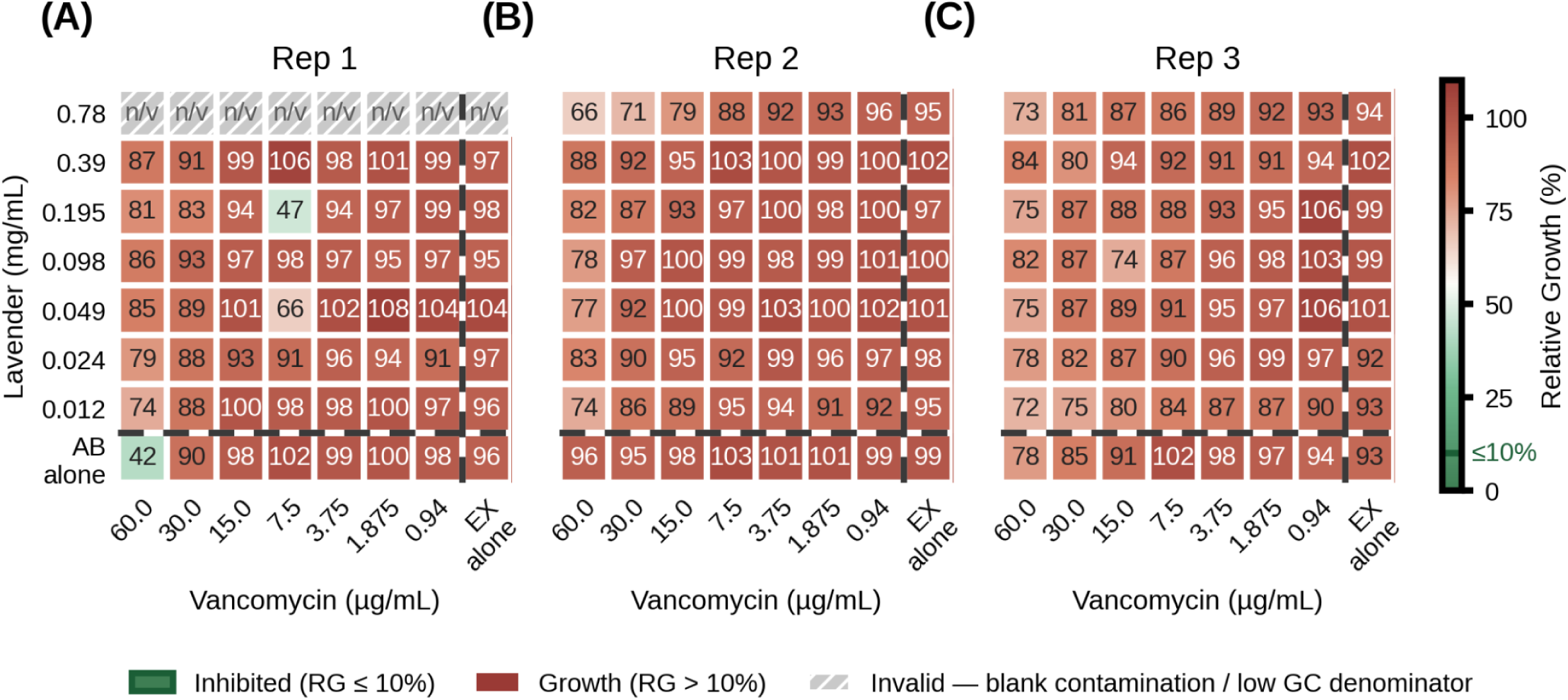
Relative growth (%) heatmaps for lavender essential oil combined with vancomycin against MRSA across three biological replicates. Rows A–G = decreasing lavender concentrations (0.78–0.012 mg/mL); column 8 = extract alone (no antibiotic). Columns 1–7 = decreasing vancomycin concentrations (60.0–0.94 µg/mL final well); Row H = antibiotic alone (no extract). Dashed lines separate the combination matrix from single-agent reference columns/rows. Green cells with bold border = inhibited (RG ≤ 10%). Grey hatched cells = invalid due to blank well contamination. Colour bar (right) is shared across all three replicates. No genuine inhibition was observed in the combination matrix. Full plate validity audit in SM7 (Tables 1 and 2).

For combination 4 (lavender + ciprofloxacin vs *E. coli* K-12), the ciprofloxacin antibiotic control demonstrated clear inhibition in replicates 1 and 3 (ΔOD 0.03–0.15 vs growth control 0.92–1.13), confirming activity; replicate 2 showed partial inhibition (ΔOD 0.12–0.29). Blank well contamination in column 12 across all three replicates required row exclusions (Figure 5 and SM7, Table 4). The lavender extract-alone column returned 82–105% RG at 1.56 mg/mL (1× MIC), the highest concentration testable within DMSO solubility constraints. No combination matrix well demonstrated inhibition consistent with combined activity beyond ciprofloxacin alone.

**Figure 3.**
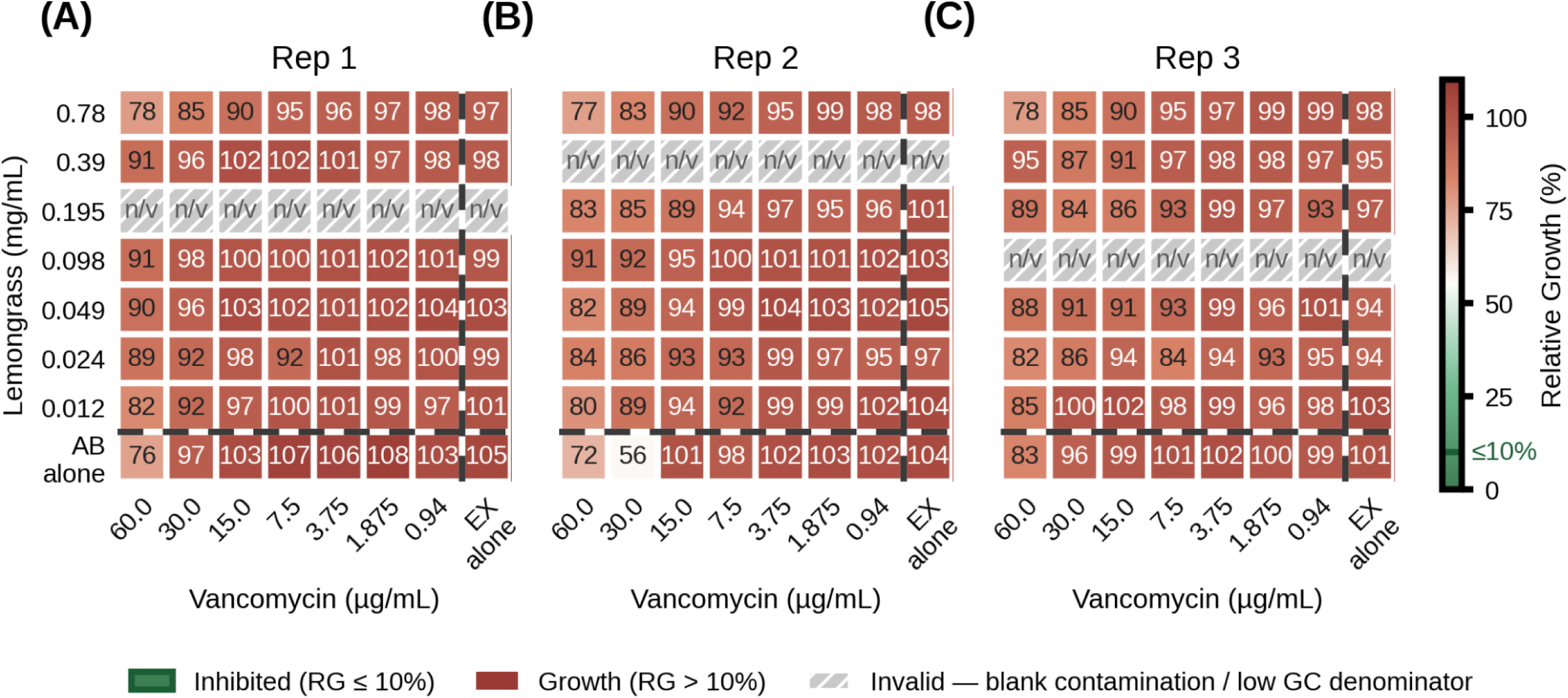
Relative growth (%) heatmaps for lemongrass essential oil combined with vancomycin against MRSA across three biological replicates. Layout as Figure 2 Rows A–G = lemongrass 0.78–0.012 mg/mL. No inhibition was observed in the combination matrix or single-agent reference rows in any replicate. Full plate validity audit in SM7 (Tables 1 and 2).

**Figure 4.**
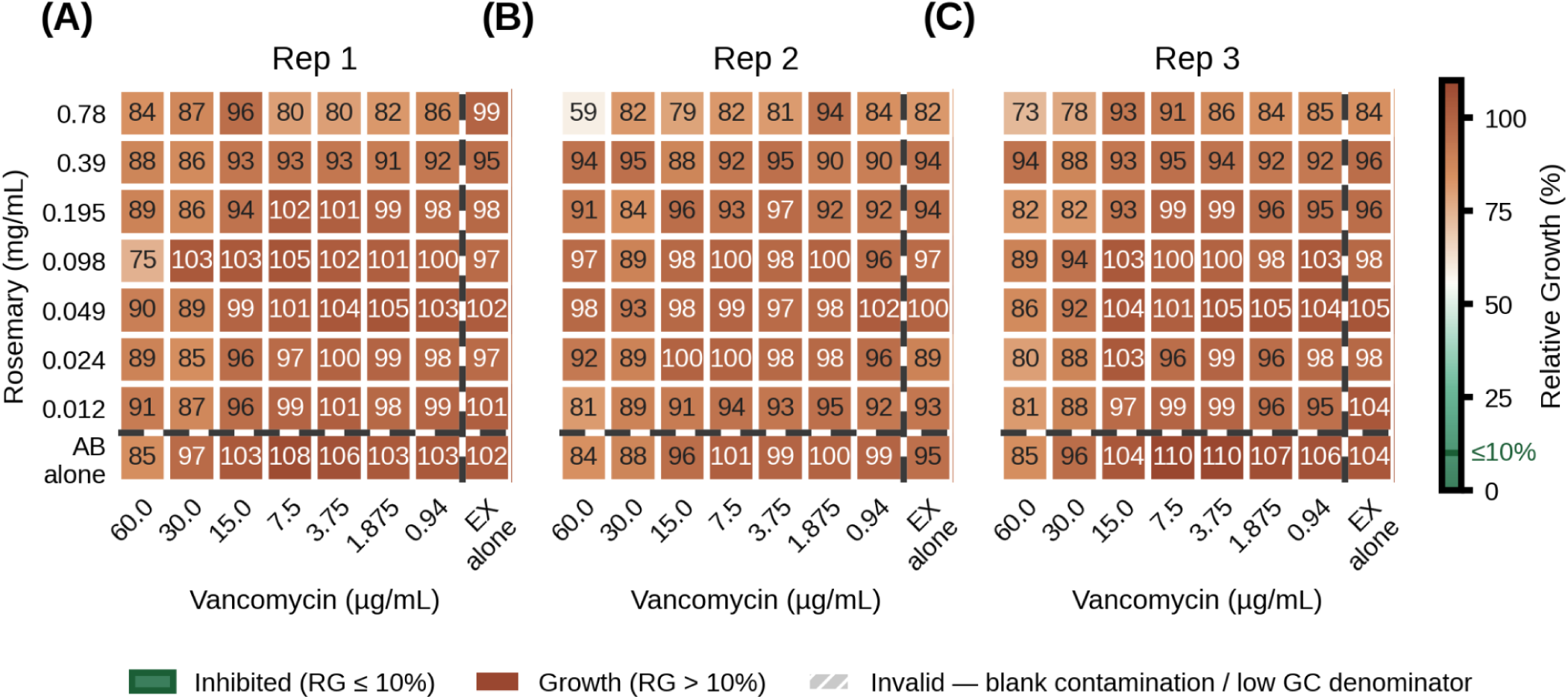
Relative growth (%) heatmaps for rosemary ethanolic extract combined with vancomycin against MSSA across three biological replicates. **(A)** Replicate 1; **(B)** Replicate 2; **(C)** Replicate 3. Layout as Figure 2: rows give rosemary concentration (0.78–0.012 mg mL⁻¹; bottom row "AB alone" = vancomycin without extract); columns give vancomycin concentration (60.0–0.94 µg mL⁻¹; right column "EX alone" = rosemary without antibiotic); dashed lines separate the combination matrix from the single-agent references. RG and the ≤ 10% inhibition criterion are as defined for Figure 2. All wells were valid (no blank-well contamination; SM7, Table 4). No inhibition was observed across the combination matrix or single-agent references in any replicate.

**Figure 5.**
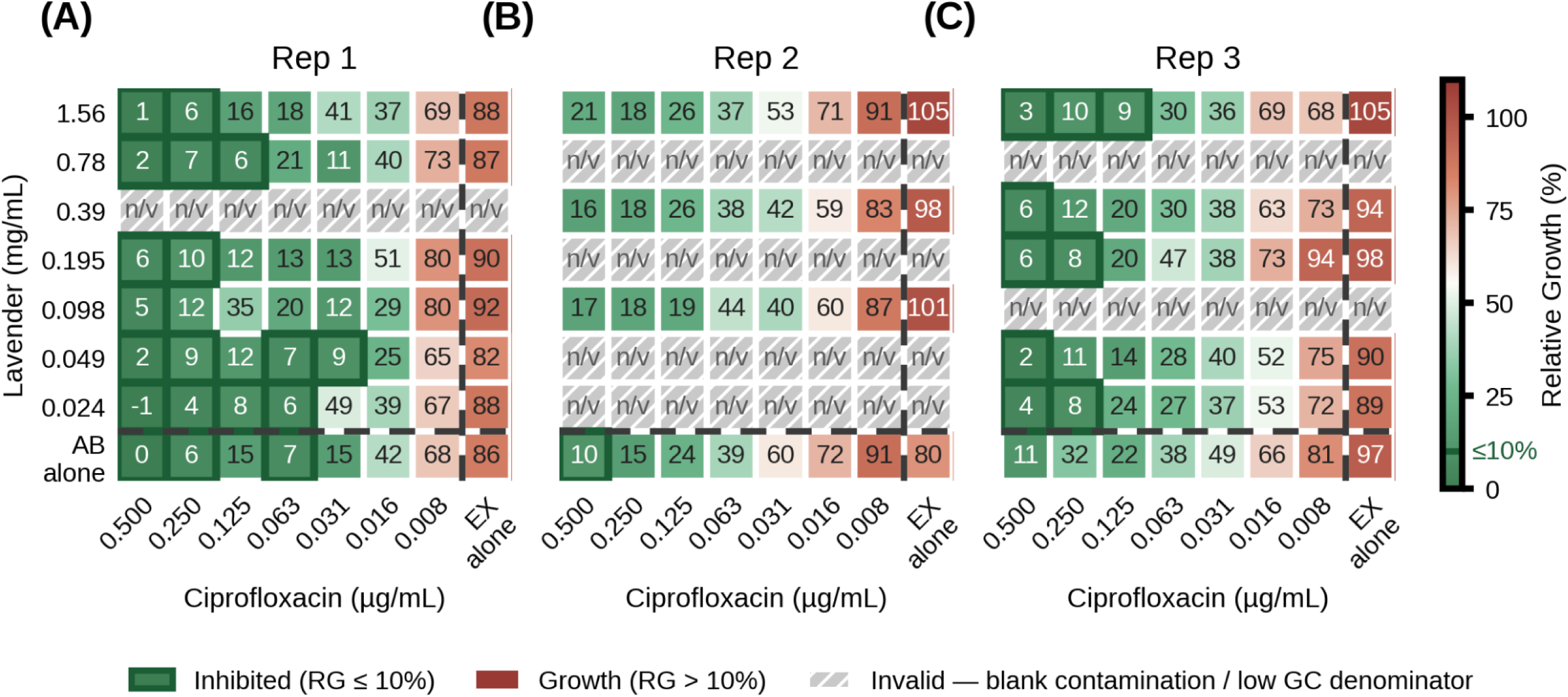
Relative growth (%) heatmaps for lavender essential oil combined with ciprofloxacin against *E. coli* K-12 across three biological replicates. **(A)** Replicate 1; **(B)** Replicate 2; **(C)** Replicate 3. Rows give lavender concentration (1.56–0.024 mg mL⁻¹; the combination began at 1× MIC owing to the 1% v/v DMSO solubility limit; bottom row "AB alone" = ciprofloxacin without extract); columns give ciprofloxacin concentration (0.500–0.008 µg mL⁻¹; right column "EX alone" = lavender without antibiotic); dashed lines separate the combination matrix from the single-agent references. RG and the ≤ 10% inhibition criterion (green, bold border) are as defined for Figure 2. Inhibition in the high-ciprofloxacin columns (0.500 and 0.250 µg mL⁻¹) appears uniformly across the lavender gradient and in the ciprofloxacin-alone row, reflecting ciprofloxacin activity at ≥ 1× MIC rather than combinatorial enhancement; no extract-alone inhibition was observed at 1× MIC. Hatched "n/v" cells were excluded owing to blank-well contamination (Replicate 1 row C; Replicate 2 rows B, D, F, G; Replicate 3 rows B, E; SM7, Table 4).

### 3.6 Chemical characterisation of plant preparations

Phytochemical screening and GC-MS profiling characterised the six preparations. Screening was restricted to ethanolic extracts; essential oil composition was determined by GC-MS. Results are summarised in Table 6, Table 1, and Figure 6.

**Figure 6.**
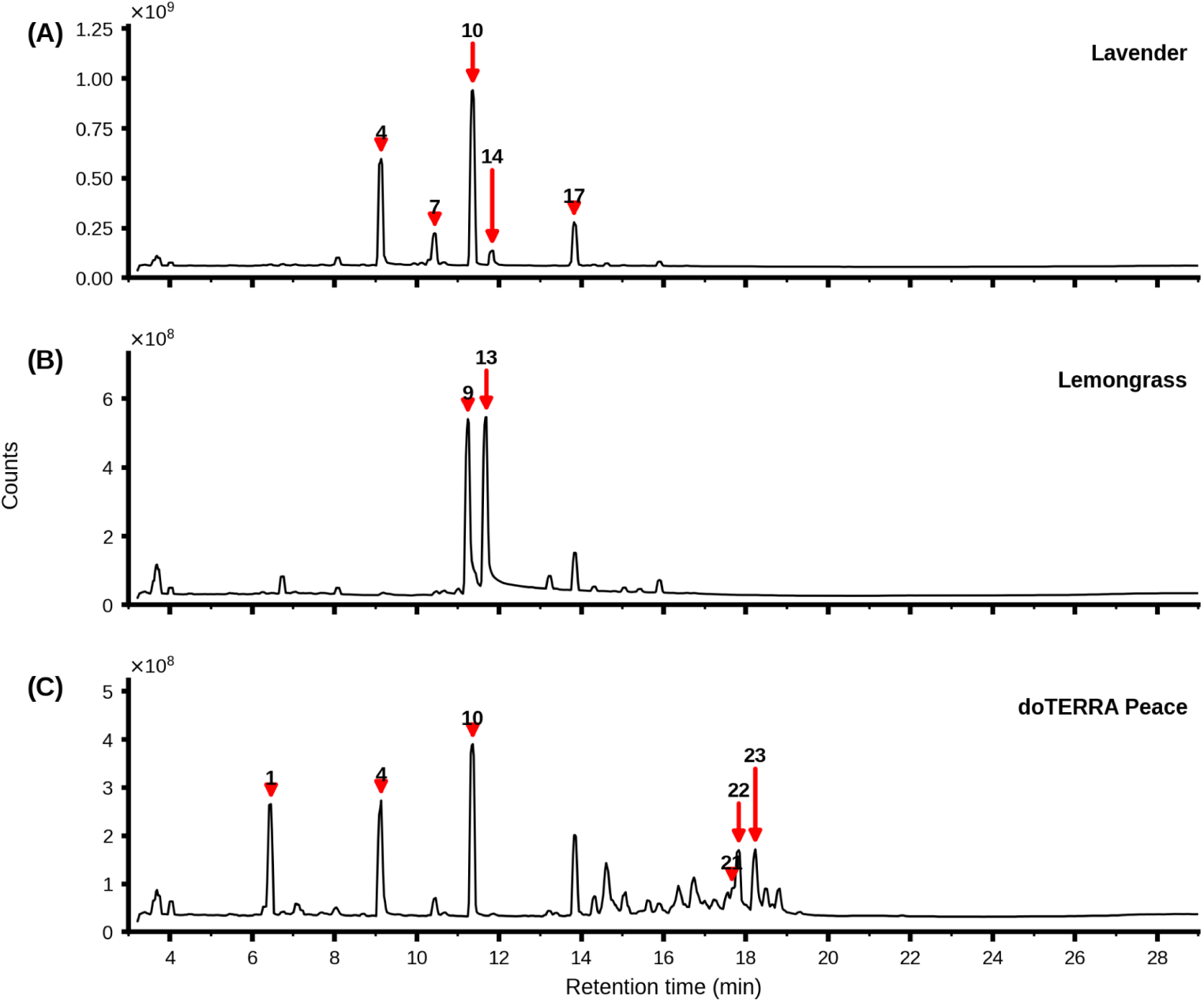
Annotated total ion chromatograms (TIC) of the three essential oil preparations at 1 mg/mL in DMSO. **(A)** lavender essential oil; **(B)** lemongrass essential oil; **(C)** doTERRA Peace blend essential oil. X-axis: retention time (min); Y-axis: total ion current (counts), each panel shown on its own intensity scale. Numbered red markers indicate the principal constituents discussed in the text; peak numbers correspond to the compound identifications and relative percentages reported in Table 1. Only major peaks were annotated for clarity. The three ethanolic extracts (nettle, thyme, rosemary) are not shown, as direct-injection GC-MS resolved no botanical terpene constituents above matrix background (Section 3.6); their qualitative composition is reported in Table 6.

**Table 6.** Qualitative phytochemical screening of ethanolic extracts of nettle, thyme, and rosemary.

| Phytochemical class | Test used | Positive indicator | Nettle | Thyme | Rosemary |
| --- | --- | --- | --- | --- | --- |
| Flavonoids | Shinoda test | Pink–crimson coloration | +(yellow–orange) | +(orange–red) | – |
| Alkaloids | Wagner's reagent | Brown precipitate | + | + | + |
| Phenolics | Ferric chloride test | Blue-black or green coloration | +(green) | +(green) | +(green) |
| Saponins | Froth test | Stable foam >1 cm | – | – | – |

+ = positive result; – = negative result. All tests conducted in triplicate using freshly prepared extracts. Assay protocol stated in SM3 (Subsections 1-4)

Phenolic compounds and alkaloids were detected in all three ethanolic extracts. Flavonoids were detected in nettle and thyme, yet absent in rosemary. Saponins were absent in all three ethanolic extracts (Table 6).

GC-MS profiling of three essential oils produced clean, interpretable profiles (Figure 6, Table 1). Lavender was a linalool chemotype, with linalyl acetate (36.1%, HIT 954) and linalool (24.8%, HIT 949) dominant. Secondary components included terpinen-4-ol (7.6%), β-caryophyllene (7.9%), and lavandulyl acetate (5.8%). Lemongrass was a citral chemotype, with geranial (43.3%, HIT 944) and neral (32.6%, HIT 894) accounting for 76% of the profile. The doTERRA Peace blend displayed the most complex profile, with 53 detected compounds including linalyl acetate (14.1%), α-pinene (6.5%), and a sesquiterpene fraction (8.4% nerolidol, RT 15–19 min).

### 3.7 Chemotype Similarity Index Analysis of Essential Oil Compositions

#### 3.7.1 Pairwise compositional similarity

The three essential oils characterised by GC-MS (Table 1) presented compositionally divergent profiles. Pairwise Pearson correlation across the union of the top 10 compounds from each preparation (n = 23 unique compounds) is presented in Table 7.

**Table 7.** Pairwise compositional similarity between essential oil preparations.

| Pair | Pearson $r$ | Interpretation |
| --- | --- | --- |
| Lavender vs Lemongrass | −0.153 | Compositionally unrelated |
| Lavender vs doTERRA Peace | +0.716 | Strong compositional similarity |
| Lemongrass vs doTERRA Peace | −0.256 | Compositionally unrelated |
| **Mean CSI (panel)** | **+0.102** | **Net compositional independence** |

The strong positive correlation between lavender and the doTERRA Peace blend (r = +0.716) reflected substantial shared dominance of linalyl acetate (36.1% in lavender; 14.1% in doTERRA) and linalool (24.8% in lavender; 7.3% in doTERRA), the two largest contributors to the cross-product sum. The two near-zero correlations of pairwise comparisons with lemongrass reflected the strict compositional independence of the citral chemotype from the linalool/linalyl acetate-dominated preparations. The mean CSI of +0.102 indicated that, taken as a panel, the three essential oils are compositionally independent overall, with the strong lavender–doTERRA Peace correspondence balanced by the divergence of lemongrass from both (Table 7).

#### 3.7.2 Strain-dependent chemistry–activity relationships

When pairwise compositional correlations were compared to the log₂-MIC differences between the same pairs of preparations against each strain (Table 8 and Figure 7), the relationship varied dramatically.

**Figure 7.**
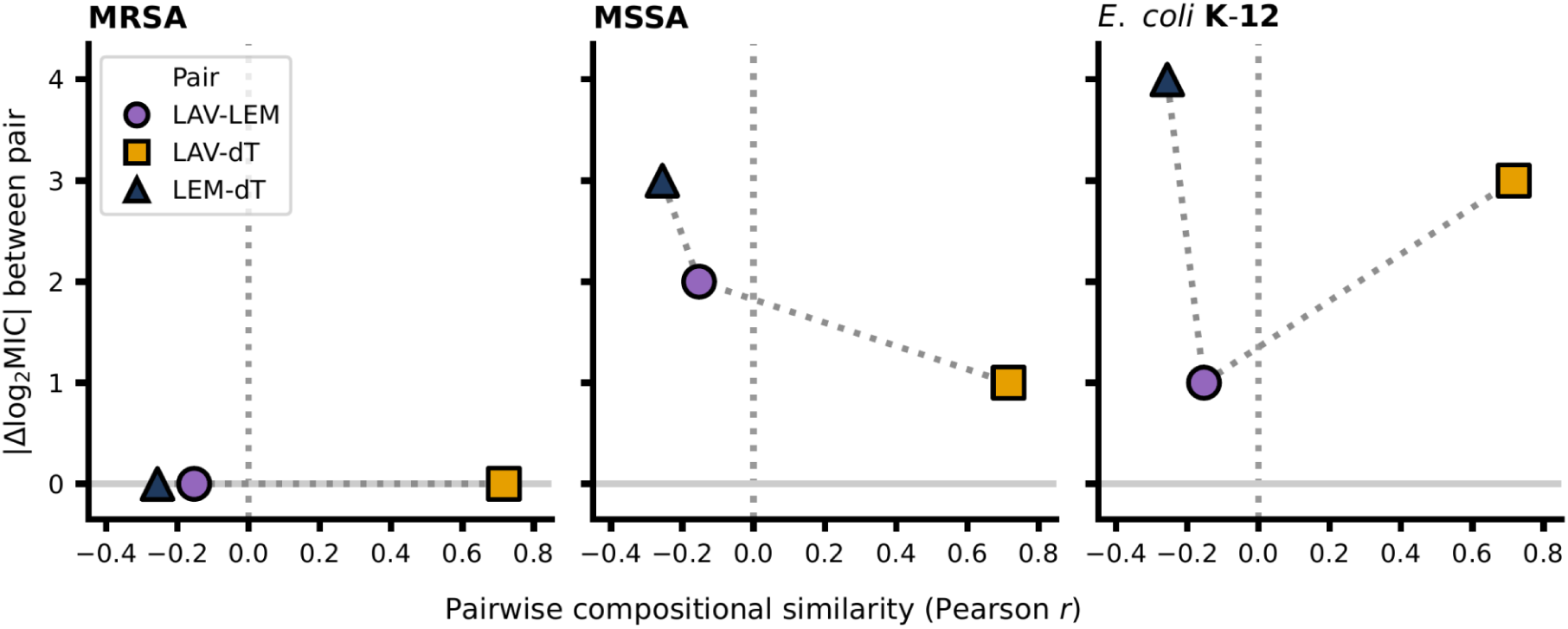
Strain-dependent relationship between essential oil compositional similarity and antimicrobial outcome similarity. Pairwise compositional similarity (Pearson *r*, across the 23-compound CSI space) plotted against absolute log₂-MIC difference (|Δlog₂MIC|) for the three essential oil pairs across the three bacterial strains: **(A)** MRSA, **(B)** MSSA, **(C)** *Escherichia coli* K-12. Pairs are identified by marker: LAV–LEM (purple circle), LAV–dT (orange square), LEM–dT (dark navy triangle). Each marker represented |Δlog₂MIC| computed from median MIC values across three biological replicates per condition; per-replicate values are reported in SM2, Table 4. Against MRSA, all three pairs converged at |Δlog₂MIC| = 0 regardless of compositional similarity, indicating that chemistry does not predict potency. Against MSSA, higher compositional similarity associated with smaller MIC differences, consistent with a monotonic chemistry–activity correspondence. Against *E. coli* K-12, compositional similarity did not track MIC differences (the most similar pair, LAV–dT, differed by 3 log₂ steps; the least similar, LEM–dT, by 4) attributable to compositional differences in the sesquiterpene fraction (Section 4.6). Dotted lines connect the three data points within each panel for visual clarity; they do not represent fitted regressions. The dotted vertical line at *r* = 0 marks the boundary between negative and positive compositional correlation; the horizontal line at |Δlog₂MIC| = 0 marks identical antimicrobial potency. *n* = 3 essential oil pairs per strain; descriptive analysis only.

**Table 8.** Pairwise compositional similarity (Pearson r) and absolute log₂-MIC differences (|Δlog₂MIC|) between essential oil pairs across the three bacterial strains tested. MIC values from Section 3.3.

| Pair | Chem. $r$ | MRSA<br>$\vert \Delta\log_2\text{MIC}\vert $ | MSSA<br>$\vert \Delta\log_2\text{MIC}\vert $ | *E. coli*<br>$\vert \Delta\log_2\text{MIC}\vert $ |
| --- | --- | --- | --- | --- |
| LAV-LEM | −0.153 | 0.00 | 2.00 | 1.00 |
| LAV-dT | +0.716 | 0.00 | 1.00 | 3.00 |
| LEM-dT | −0.256 | 0.00 | 3.00 | 4.00 |

Against MRSA, all three essential oils produced an identical median MIC of 0.39 mg/mL (Section 3.3) despite spanning the full range of pairwise compositional similarity (*r* = −0.256 to +0.716). The log₂-MIC difference was zero across all three pairs (Table 8). Compositional similarity did not predict anti-MRSA potency in this dataset.

Against MSSA, log₂-MIC differences spanned 1.0 to 3.0 across the three pairs. The pair with strongest compositional similarity (LAV–dT, *r* = +0.716) showed the smallest log₂-MIC difference (Δlog₂ = 1.0; lavender 1.56 mg/mL vs doTERRA 3.13 mg/mL). The pair with weakest compositional similarity (LEM–dT, *r* = −0.256) showed the largest log₂-MIC difference (Δlog₂ = 3.0; lemongrass 0.39 mg/mL vs doTERRA 3.13 mg/mL). Compositional similarity was broadly consistent with antimicrobial similarity against MSSA, describing a monotonic relationship.

Against *E. coli* K-12, the monotonic relationship broke down. The most compositionally similar pair (LAV–dT, r = +0.716) differed by Δlog₂ = 3.0 (lavender 1.56 mg/mL vs doTERRA 12.5 mg/mL), while the least similar pair (LEM–dT, r = −0.256) showed the largest difference (Δlog₂ = 4.0; lemongrass 0.78 mg/mL vs doTERRA 12.5 mg/mL). Compositional similarity therefore did not predict antimicrobial similarity against *E. coli*.

The strain-dependent chemistry–activity relationship — convergent against MRSA, monotonic against MSSA, and absent against E. coli — was the principal CSI finding of this study (Figure 7).

## 4. DISCUSSION

The findings considered here addressed that firstly, the preparations would show measurable inhibitory activity and antibiotic interactions classifiable by FICI; and secondly, that compositional similarity between essential oils would relate to antimicrobial activity in a strain-dependent manner. The former was partially supported — reproducible inhibitory activity was confirmed by broth microdilution, but no combination produced a calculable FICI, all being classified as indifference. The latter was supported and refined: compositional similarity related to activity, but the strength and direction of that relationship varied across strains.

### 4.1 The disc diffusion–broth microdilution discrepancy and its methodological implications

The most striking observation was the failure of disc diffusion to detect activity for lemongrass despite its lowest median MIC values (0.39 mg/mL against both MRSA and MSSA). All six extract preparations failed to produce zones of inhibition, despite five preparations subsequently demonstrating reproducible MIC values at or below 3.13 mg/mL by broth microdilution. This is attributable to the physicochemical properties of the preparations: lipophilic essential oils and crude ethanolic extracts in DMSO diffuse poorly through agar, reducing assay sensitivity relative to water-soluble antibiotics for which the method was originally developed (Balouiri *et al*., 2016). Antibiotic control performance under identical conditions confirmed failure was preparation-specific. Diffusion-based screening cannot serve as a primary or exclusionary method for plant-derived preparations; broth microdilution should be the default (Balouiri *et al*., 2016).

### 4.2 Convergent anti-MRSA potency across compositionally distinct essential oils

Three essential oils achieved an identical median MIC of 0.39 mg/mL against MRSA — lavender, lemongrass, and doTERRA Peace — despite distinct chemical compositions.

Lavender was dominated by linalyl acetate and linalool (61% combined; linalool chemotype). Lemongrass was dominated by geranial and neral (76% combined; citral chemotype). The doTERRA blend was a sesquiterpene-rich mixture with linalyl acetate at only 14.1%.

Comparable convergent potency has been reported: Oliveira *et al*., (2021) reported citral MIC values of 5–40 mg/mL against MRSA clinical isolates, and Cavanagh and Wilkinson (2005) reported linalool-rich lavender preparations active in the 0.5–4 mg/mL range. The 0.39 mg/mL convergence here sits at the potent end of these ranges.

The most parsimonious interpretation is that all three preparations acted via membrane-targeted disruption of the Staphylococcal cell envelope through chemically distinct routes converging on a similar functional outcome. Linalool and linalyl acetate increase membrane permeability and disrupt the proton motive force in *S. aureus* (Raut and Karuppayil, 2014). Citral and geranial also act via membrane disruption but additionally interfere with ATP synthesis and quorum-sensing pathways (Oliveira *et al*., 2021; Khwaza and Aderibigbe, 2025). The doTERRA blend’s mixed monoterpene–sesquiterpene fraction matched the potency of the single-origin oils despite different dominant chemistries. This suggested potency may be governed less by the dominant constituent than by cumulative lipophilic terpene partitioning — a hypothesis warranting confirmation by membrane permeability and ATP leakage assays.

This aligns with literature reporting structurally dissimilar terpene mixtures producing similar antibacterial outcomes (Bakkali *et al*., 2008; Nazzaro *et al*., 2013), suggesting that membrane perturbation in *S. aureus* may be governed by an aggregate property of the lipophilic constituent fraction rather than by single-compound selectivity. This convergence was quantified using the Chemotype Similarity Index (Section 4.6), which formalised the observation that compositional similarity does not predict anti-MRSA potency in this panel.

Reinforcing this interpretation is the divergence in compositional complexity across the three preparations. The doTERRA Peace blend contained 53 detected compounds across multiple structural classes, whereas lavender was dominated by only two compounds (linalyl acetate and linalool, 61% combined) and lemongrass by two (geranial and neral, 76% combined).

Preparations differing widely in compositional complexity — from the sesquiterpene-rich doTERRA blend to the two-compound-dominated lavender and lemongrass oils — converged on identical anti-MRSA potency, suggesting that complexity itself was not a primary determinant of activity in this dataset, with cumulative lipophilic load and dominant compound classes appearing to drive the observed convergence.

### 4.3 The Gram-negative permeability barrier and differential susceptibility

Differential susceptibility between Gram-positive and Gram-negative microorganisms was most evident for the doTERRA blend (MRSA MIC 0.39 mg/mL versus *E. coli* MIC 12.5 mg/mL — a 32-fold difference). The single-origin essential oils showed smaller shifts in the same direction. The mechanism is well established: the Gram-negative outer membrane restricts penetration of lipophilic terpenes (Nazzaro *et al*., 2013). The 32-fold reduction for the doTERRA blend, contrasted with the 4-fold reduction for lavender (0.39 to 1.56 mg/mL), was interpretable from GC-MS compositions. The doTERRA blend contained a large oxygenated sesquiterpene fraction (8.4% nerolidol alone, plus cadinene-type and aromadendrene-type sesquiterpenes accounting for over 8% combined). Sesquiterpenes are larger, more lipophilic, and less effective at penetrating the outer membrane than the monoterpenes that dominated lavender (Nazzaro *et al*., 2013). The differential therefore reflected compositional differences rather than a uniform terpene effect. All three ethanolic extracts retained measurable activity against E. coli (1.56–3.13 mg/mL), comparable to or better than the doTERRA blend — consistent with phenolic disruption of outer membrane integrity through a mechanism distinct from terpene partitioning (Cowan, 1999).

Compositional complexity therefore appeared strain-dependent: neutral against Gram-positive cell envelopes where total terpene load drives outcome, but a liability against Gram-negative outer membrane filtering where larger-molecular-weight compounds penetrated poorly. The chemometric consequences of this compositional difference — specifically the failure of whole-profile similarity to capture size-class-dependent penetration — are developed in Section 4.6.

### 4.4 Bacteriostatic rather than bactericidal activity at tested concentrations

The post-exposure culturability data resolved a question MIC data cannot address: whether inhibitory activity translated into killing. No preparation produced a ≥3 log₁₀ CFU mL⁻¹ reduction at any timepoint (Wiegand *et al*., 2008). The most pronounced reductions — lavender 2× MIC against MRSA achieving 2.43 log₁₀ at 2 hours — were transient, with counts recovering to near-baseline by 24 hours. This was consistent with bacteriostatic rather than bactericidal activity at concentrations achievable within DMSO solubility constraints.

The recovery to baseline carries clinical implications: therapeutic application would require formulation strategies supporting continuous exposure rather than a single bactericidal dose. The interpretive limitation imposed by the absence of a validated neutralisation step is acknowledged.

### 4.5 Indifference in checkerboard combinations and the inoculum effect

All four checkerboard combinations were classified as indifference, consistent with the literature on essential oil–antibiotic interactions, which reported indifference as the predominant outcome for terpene-rich preparations combined with glycopeptide and fluoroquinolone antibiotics (Langeveld *et al*., 2014; Soulaimani, 2025). Variable interaction outcomes including indifference have also been reported for citral-rich preparations combined with conventional antibiotics against MRSA (Oliveira *et al.,* 2021). In combinations 1–3, the vancomycin-alone reference failed to demonstrate inhibition at 2× MIC despite the standalone MIC being well-characterised in the same media. This was most plausibly attributable to an inoculum-mediated upward shift in vancomycin MIC where elevated inoculum density increased the drug requirement for inhibition (Wiegand *et al*., 2008). The use of BHI broth rather than cation-adjusted Mueller-Hinton broth (CAMHB) likely compounded this; BHI’s higher protein content can elevate effective vancomycin MIC by binding free drug (Van de Vel *et al*., 2019). For combination 4, ciprofloxacin remained active, but the maximum lavender concentration testable within the 1% DMSO threshold was 1× MIC, limiting the matrix range over which synergy could be detected. The indifference classification was methodologically constrained and does not preclude synergy at higher lavender concentrations or under alternative solvent conditions, which remain open questions for future work.

### 4.6 Chemotype Similarity Index reveals strain-dependent chemistry–activity relationships

The Chemotype Similarity Index (CSI) was introduced to convert the qualitative observation of convergent potency (Section 4.2) into a quantitative relationship between compositional similarity and antimicrobial outcome. Applied across the three essential oils, it revealed that this relationship was not fixed but varied in both strength and direction across the three strains.

Against MRSA, the three oils achieved an identical median MIC (0.39 mg/mL) despite spanning a Pearson *r* range of nearly one full unit (−0.256 to +0.716); the most and least compositionally similar pairs produced the same MIC outcome. Composition was therefore informative for identifying an oil but uninformative for predicting its anti-MRSA potency. Against MSSA the relationship became monotonic — the strong-similarity pair (LAV–dT) showed the smallest activity difference (Δlog₂ = 1.0) and the weakest-similarity pair (LEM–dT) the largest (Δlog₂ = 3.0) — indicating that compositional similarity was a partial, direction-aligned predictor against the sensitive strain. Against *E. coli* the pattern broke down; LAV–dT differed by 3.0 despite being the most similar, while the least similar LEM–dT differed most (4.0). This breakdown was mechanistically interpretable — lavender and the doTERRA blend shared dominant monoterpenes but differed in the oxygenated sesquiterpene fraction (Section 4.3), and against a microorganism whose primary defence is permeability filtering, that functionally decisive difference was invisible to a whole-profile correlation. Pearson similarity does not distinguish freely permeating monoterpenes from size-restricted sesquiterpenes, and against a Gram-negative target this compositional blindness produced the observed breakdown. The divergence between the MRSA convergence and the MSSA gradient was also notable: methicillin resistance (*mecA*/PBP2a) alters cell-wall biosynthesis rather than membrane permeability directly (Chambers and DeLeo, 2009; Lakhundi and Zhang, 2018), therefore the flattened MIC landscape against MRSA is a hypothesis for direct tests by membrane-permeability and ATP-leakage assays across matched MRSA/MSSA pairs, rather than an established effect.

These trends must be read against the scale of the dataset. The CSI analysis was exploratory and descriptive: three preparations cannot support inferential testing of the correlation coefficients, and no p-values are claimed. The framework was presented as proof-of-concept for application to larger panels (n ≥ 20 essential oils), where the trends suggested herein could be tested with adequate power. The three ethanolic extracts were excluded because direct-injection GC-MS without solid-phase cleanup could not resolve their botanical terpene profiles (Myers et al., 2021), restricting current applicability to volatile-rich preparations; solid-phase extraction with LC-MS profiling would extend CSI to aqueous and ethanolic extracts.

Compositional-similarity frameworks are well established in chemometric authentication of plant materials (Adams, 2007), but their integration with quantitative antimicrobial outcomes remains rare; most plant-antimicrobial studies report composition without relating it analytically to MIC (Marchese et al., 2016). CSI addresses that gap as a quantitative bridge between the two datasets. The central methodological implication is that such a framework must be strain-aware: a single similarity–activity correlation pooled across microorganisms would obscure the reversal documented here and yield misleadingly low predictive value.

CSI is therefore a framework rather than a finished tool — its contribution in this study was to reframe the convergent MRSA observation quantitatively and to provide a reusable scaffold that larger panels and matched strain pairs can populate. The strain-dependent pattern itself — convergent against MRSA, monotonic against MSSA, absent against E. coli — is a novel observation in the plant-antimicrobial literature and warrants validation across larger essential oil panels and additional Gram-negative strains.

### 4.7 Further Work

The findings of this study identified five specific methodological and mechanistic directions for future research, each addressing a defined limitation as discussed above.

The post-exposure culturability assay should be repeated with a validated antimicrobial neutralisation step (Dey-Engley broth or equivalent) and multiple biological replicates, enabling formal time-kill kinetic analysis and reliable distinction between bacteriostatic and bactericidal effects across the preparation panel.

The inoculum-mediated upward shift in vancomycin MIC observed in checkerboard conditions warrants systematic evaluation across CAMHB and BHI media using European Committee on Antimicrobial Susceptibility Testing defined inoculum standards, to determine whether the indifference outcome reflects a genuine pharmacological interaction or a nutrient-mediated artefact.

The laboratory-maintained MRSA isolate should be replaced with a characterised reference strain such as ATCC 33591, with whole-genome sequencing performed to confirm resistance gene complement and exclude potential phenotypic drift arising from extended subculture.

GC-MS profiles of the three ethanolic extracts could not be resolved by direct injection due to matrix interference; future characterisation of nettle, thyme, and rosemary preparations should employ solid-phase extraction cleanup or LC-MS to obtain interpretable compositional data, enabling chemical comparison with the essential oils.

The convergent anti-MRSA activity (MIC 0.39 mg/mL) observed across three compositionally divergent essential oils — lavender, lemongrass, and the doTERRA Peace blend — suggests a shared membrane-disrupting mechanism that should be confirmed directly by membrane permeability assays, ATP leakage measurements, and quantification of cellular potassium efflux.

### 4.8 Conclusion

The principal aim — to evaluate the antimicrobial activity, synergistic potential, and phytochemical composition of selected medicinal plant extracts and essential oils against clinically relevant bacterial pathogens — was achieved. The first objective, to determine inhibitory activity and characterise antibiotic interactions, was partially met: reproducible inhibitory activity was confirmed by broth microdilution, with three essential oils achieving 0.39 mg/mL against MRSA and rosemary achieving the same against MSSA, but no combination yielded a numerically calculable FICI. All four combinations were instead classified as indifference under the Odds (2003) framework, with inhibition boundaries unresolvable in the reference controls — an outcome attributable to inoculum-mediated and solubility-related methodological constraints rather than to a true absence of interaction. The second objective, to relate compositional similarity between essential oils to antimicrobial activity, was met and refined: similarity related to activity, but the strength and direction of that relationship reversed across strains.

The convergent anti-MRSA potency across three compositionally divergent essential oils was the most biologically significant finding and warrants direct mechanistic investigation. The Chemotype Similarity Index introduced here reframes this observation quantitatively: across a Pearson r range spanning nearly a full unit, compositional similarity did not predict anti-MRSA potency, supporting a shared membrane-targeted mechanism operating through chemically distinct routes. The principal methodological contribution of this study is the strain-dependent chemistry–activity relationship that CSI exposes — convergent against MRSA, monotonic against MSSA, and absent against E. coli K-12 — which indicated that any compositional framework intended to predict antimicrobial activity must be applied on a strain-specific basis. While derived from only three essential oil preparations and therefore exploratory, this pattern is a novel observation in the plant antimicrobial chemometric literature and provides a reusable analytical scaffold for validation across larger preparation panels.

The transient sub-bactericidal pattern indicated that any clinical translation would require formulation strategies supporting continuous exposure. Taken together, these findings support the case for plant-derived preparations as adjuncts rather than replacements for conventional antibiotics, and rest on the specific mechanistic and methodological refinements set out in Section 4.7.

## Supporting information

Supplementary_Material.docx

## Author Contributions

**AB:** Conceptualization, Methodology, Investigation, Formal analysis, Data curation, Software, Visualization, Writing – original draft, Writing – review & editing. **AS:** Conceptualization, Supervision, Writing – review & editing. Both authors contributed to the article and approved the submitted version.

## Acknowledgments

The authors would like to thank Dr Samantha Bowerbank (School of Geography and Natural Sciences, Northumbria University) for performing the gas chromatography–mass spectrometry profiling of the essential oil preparations.

The Chemotype Similarity Index framework and analytical approach were conceived and developed by the lead author (AB), and the data analysis was performed by the lead author. A generative AI tool (Anthropic Claude, Claude Opus 4.5; Anthropic, https://claude.ai) was used to assist with implementing and verifying the computational analysis in Python, and with generating the publication figures from the authors’ own underlying data. All AI-assisted outputs, including figures, were checked by the authors against the source data for accuracy. All conceptual contributions, data, interpretation, and conclusions are the authors’ own, and the authors take full responsibility for the accuracy of all reported results, including all AI-assisted content.

## Data Availability Statement

The Chemotype Similarity Index calculation code and example input data analysed in this study are openly available so that the framework may be applied to other compositional datasets, at: https://github.com/atulbhat080-crypto/CSI-Framework and archived with a permanent identifier at https://doi.org/10.5281/zenodo.20725211. All other data generated and analysed during this study are included in this article and its Supplementary Material.

