## Supplementary_Material.docx for "Convergent anti-MRSA potency across compositionally distinct essential oils: a chemotype similarity index for strain-dependent chemistry–activity analysis"

The following supplementary materials provide per-replicate raw and summary data, plate validity audits, worked calculations, and reagent protocols supporting the main manuscript. SM1 contains disc diffusion zone measurements. SM2 contains per-replicate plant-preparation MIC values across all biological replicates, with one representative plate shown in full plate-level detail. SM3 and SM4 provide the phytochemical screening protocols and GC–MS interpretability notes, and the worked Chemotype Similarity Index calculation, supporting the chemometric framework. SM5 and SM6 provide the post-exposure culturability and antibiotic MIC data respectively, and SM7 provides the checkerboard FICI synergy assay data with its plate validity audit.

**Contents**

Supplementary Material (SM) 1 — Disc Diffusion Zone Measurements

Supplementary Material (SM) 2 — Plant Preparation MIC Determination

Supplementary Material (SM) 3 — Phytochemical Screening Protocols and Notes on GC–MS Interpretability

Supplementary Material (SM) 4 — Worked Chemotype Similarity Index Calculation

Supplementary Material (SM) 5 — Post-Exposure Culturability

Supplementary Material (SM) 6 — Antibiotic MIC Determination

Supplementary Material (SM) 7 — Checkerboard FICI Synergy Assay

**Supplementary Material 1**

**Disc Diffusion Zone Measurements**

Per-replicate zone of inhibition measurements supporting Section 3.2 and Table 2 in the main manuscript. Zones were recorded at the bench as zone radius in centimetres on the underside of each agar plate using a calibrated ruler, then converted to total zone diameter in millimetres and zone diameter beyond the 6 mm filter paper disc edge. All measurements are n = 3 independent biological replicates per preparation–strain combination. Lemongrass essential oil produced zone radii of 0.30 cm (i.e., disc edge only, no zone) in all three replicates against all three strains and is reported as 0.0 ± 0.0 mm.

**1. Measurement Convention**

Zones were recorded as zone radius in centimetres (radius from the centre of the disc to the outer edge of the inhibition zone). Each radius was converted in two steps:

*Step 1 — total zone diameter (mm) = radius (cm) × 2 × 10*

*Step 2 — zone beyond disc edge (mm) = total zone diameter (mm) − 6 mm (disc diameter)*

Zone diameters beyond the 6 mm disc edge are the values reported in main manuscript Section 3.2 and Table 1, and are used for inter-preparation comparison. A radius reading of 0.30 cm (= total zone diameter 6 mm = zone beyond disc 0 mm) indicates no inhibition beyond the disc edge.

**2. Per-Replicate Measurements**

**Table 1. Per-replicate disc diffusion measurements: raw radius readings (cm), computed total zone diameters (mm), and zone diameters beyond the 6 mm disc edge (mm).**

*All conversions follow the formula in Subsection 1. Means and standard deviations are reported for the zone-beyond-disc values; SDs calculated as sample SD (n−1 denominator).*

| **Strain** | **Sample** | **R1 (cm)** | **R2 (cm)** | **R3 (cm)** | **R1 diam (mm)** | **R2 diam (mm)** | **R3 diam (mm)** | **R1 zone (mm)** | **R2 zone (mm)** | **R3 zone (mm)** | **Mean ± SD (mm)** |
| --- | --- | --- | --- | --- | --- | --- | --- | --- | --- | --- | --- |
| **MRSA** | Vancomycin (30 µg, positive) | 0.50 | 0.55 | 0.55 | 10 | 11 | 11 | 4 | 5 | 5 | 4.7 ± 0.6 |
|  | DMSO (1% v/v, negative) | 0.35 | 0.50 | 0.30 | 7 | 10 | 6 | 1 | 4 | 0 | 1.7 ± 2.1 |
|  | Nettle EE (50 mg/mL) | 0.30 | 0.50 | 0.30 | 6 | 10 | 6 | 0 | 4 | 0 | 1.3 ± 2.3 |
|  | Nettle EE (25 mg/mL) | 0.30 | 0.30 | 0.30 | 6 | 6 | 6 | 0 | 0 | 0 | 0.0 ± 0.0 |
|  | Lavender EO (10% w/v) | 0.45 | 0.40 | 0.30 | 9 | 8 | 6 | 3 | 2 | 0 | 1.7 ± 1.5 |
|  | Lemongrass EO (10% w/v) | 0.30 | 0.30 | 0.30 | 6 | 6 | 6 | 0 | 0 | 0 | 0.0 ± 0.0 |
|  | doTERRA Peace (10% w/v) | 0.70 | 0.50 | 0.50 | 14 | 10 | 10 | 8 | 4 | 4 | 5.3 ± 2.3 |
| **MSSA** | Vancomycin (30 µg, positive) | 0.50 | 0.55 | 0.50 | 10 | 11 | 10 | 4 | 5 | 4 | 4.3 ± 0.6 |
|  | DMSO (1% v/v, negative) | 0.30 | 0.35 | 0.30 | 6 | 7 | 6 | 0 | 1 | 0 | 0.3 ± 0.6 |
|  | Nettle EE (50 mg/mL) | 0.35 | 0.30 | 0.30 | 7 | 6 | 6 | 1 | 0 | 0 | 0.3 ± 0.6 |
|  | Thyme EE (50 mg/mL) | 0.30 | 0.30 | 0.30 | 6 | 6 | 6 | 0 | 0 | 0 | 0.0 ± 0.0 |
|  | Thyme EE (25 mg/mL) | 0.35 | 0.30 | 0.30 | 7 | 6 | 6 | 1 | 0 | 0 | 0.3 ± 0.6 |
|  | Rosemary EE (50 mg/mL) | 0.30 | 0.50 | 0.40 | 6 | 10 | 8 | 0 | 4 | 2 | 2.0 ± 2.0 |
|  | Lemongrass EO (10% w/v) | 0.30 | 0.30 | 0.30 | 6 | 6 | 6 | 0 | 0 | 0 | 0.0 ± 0.0 |
| ***E. coli* K-12** | Ciprofloxacin (5 µg, positive) | 0.90 | 0.90 | 0.975 | 18 | 18 | 19.5 | 12 | 12 | 13.5 | 12.5 ± 0.9 |
|  | DMSO (1% v/v, negative) | 0.30 | 0.30 | 0.30 | 6 | 6 | 6 | 0 | 0 | 0 | 0.0 ± 0.0 |
|  | Nettle EE (50 mg/mL) | 0.30 | 0.30 | 0.40 | 6 | 6 | 8 | 0 | 0 | 2 | 0.7 ± 1.2 |
|  | Rosemary EE (50 mg/mL) | 0.65 | 0.60 | 0.50 | 13 | 12 | 10 | 7 | 6 | 4 | 5.7 ± 1.5 |
|  | Lavender EO (10% w/v) | 0.30 | 0.30 | 0.60 | 6 | 6 | 12 | 0 | 0 | 6 | 2.0 ± 3.5 |
|  | Lemongrass EO (10% w/v) | 0.30 | 0.30 | 0.30 | 6 | 6 | 6 | 0 | 0 | 0 | 0.0 ± 0.0 |
|  | doTERRA Peace (10% w/v) | 0.50 | 0.30 | 0.30 | 10 | 6 | 6 | 4 | 0 | 0 | 1.3 ± 2.3 |

*EE = ethanolic extract; EO = essential oil. Mean ± SD reported for zone-beyond-disc values across the three replicates per row, with SD calculated as sample standard deviation (n−1 denominator). Mean values in the final column reproduce those reported in main manuscript Table 2.*

**3. Worked Calculation Example**

**Rosemary ethanolic extract at 50 mg/mL versus *E. coli* K-12 (largest plant zone observed in this assay):**

Step 1 — radius readings to total diameter:

• Plate 1: 0.65 cm × 2 × 10 = 13 mm

• Plate 2: 0.60 cm × 2 × 10 = 12 mm

• Plate 3: 0.50 cm × 2 × 10 = 10 mm

Step 2 — total diameter to zone beyond disc:

• Plate 1: 13 − 6 = 7 mm

• Plate 2: 12 − 6 = 6 mm

• Plate 3: 10 − 6 = 4 mm

Step 3 — mean and sample standard deviation:

• Mean = (7 + 6 + 4) ÷ 3 = 5.67 mm

• Sample SD = √[((7 − 5.67)² + (6 − 5.67)² + (4 − 5.67)²) ÷ 2] = √(2.33) = 1.53 mm

**• Reported as 5.7 ± 1.5 mm in main manuscript Section 3.2 and Table 2.**

**4. Notes on Interpretability**

No plant preparation produced a zone meeting standard interpretive breakpoints for antimicrobial susceptibility. The most reproducible plant result was rosemary against *E. coli* (5.7 ± 1.5 mm); the largest mean was doTERRA Peace against MRSA (5.3 ± 2.3 mm), but inter-replicate variability (4–8 mm) precluded reliable interpretation. Lemongrass produced no zone against any strain (0.0 ± 0.0 mm across all three) despite returning the lowest median MIC values in subsequent broth microdilution (Section 3.3 of the main manuscript), demonstrating a known limitation of disc diffusion for lipophilic essential oil preparations (Balouiri et al., 2016).

The DMSO 1% (v/v) negative control produced minor non-zero readings against MRSA (1.7 ± 2.1 mm) and MSSA (0.3 ± 0.6 mm) but no measurable inhibition against *E. coli* K-12 (0.0 ± 0.0 mm). The MRSA and MSSA readings are attributable to mechanical disturbance of the agar surface during disc placement rather than solvent antimicrobial activity, given that 1% DMSO is well below the threshold reported to affect bacterial viability and that the variability across replicates exceeds the mean in both cases.

All values reported in Section 3.2 of the main manuscript are derived directly from the per-replicate dataset shown in Table 1 using the conversion formula in Subsection 1.

**Supplementary Material 2**

**Plant Preparation MIC Determination**

Per-replicate minimum inhibitory concentration (MIC) values supporting Table 3 in the main manuscript. MIC was determined by broth microdilution in BHI broth across nine 96-well plates (three plates per strain × three biological replicates). Rep 1 was conducted in December 2025; Reps 2 and 3 were conducted in February 2026. One representative replicate (MRSA Rep 1) is shown in full plate-level detail in Subsection 3; per-replicate MIC values across all nine plates are summarised in Table 4 (Subsection 5).

**1. Relative Growth Formula**

Relative growth (RG) was calculated for every well as:

$$\mathrm{RG}\left( \% \right)=\frac{\Delta\mathrm{OD}_{\mathrm{sample}}- \Delta\mathrm{OD}_{\mathrm{blank}}}{\Delta\mathrm{OD}_{solvent control}- \Delta\mathrm{OD}_{\mathrm{blank}}}\times100$$

where ΔOD = OD₆₀₀ at T24 minus OD₆₀₀ at T0 for each well; ΔOD blank = row-specific mean of columns 9 and 12 (BHI broth, no bacteria); ΔOD solvent control = column 11 (bacteria in BHI + 1% v/v DMSO, no extract). MIC was defined as the lowest concentration at which RG ≤ 10%.

**2. Plate Layout**

**Table 3.** Each 96-well plate contained six plant preparations (rows A–F) tested in two-fold serial dilution across columns 1–8, plus shared control columns:

| **Region** | **Content** |
| --- | --- |
| Row A | Nettle ethanolic extract (*Urtica dioica*) |
| Row B | Thyme ethanolic extract (*Thymus vulgaris*) |
| Row C | Rosemary ethanolic extract (*Rosmarinus officinalis*) |
| Row D | Lavender essential oil (*Lavandula angustifolia*) |
| Row E | Lemongrass essential oil (*Cymbopogon flexuosus*) |
| Row F | doTERRA Peace blend essential oil |
| Cols 1–8 | Two-fold serial dilutions: 50, 25, 12.5, 6.25, 3.13, 1.56, 0.78, 0.39 mg/mL |
| Col 9 | Blank (BHI broth only, no bacteria) — row-specific blank correction |
| Col 10 | Additional blank reference |
| Col 11 | Solvent control (bacteria in BHI + 1% v/v DMSO, no extract) — RG denominator |
| Col 12 | Additional blank reference — mean blank calculation |

**3. Representative Plate — MRSA Rep 1**

Tables 1 and 2 show the complete 0h and 24h OD₆₀₀ readings for one representative plate (MRSA Rep 1). Equivalent plates were generated for MSSA and *E. coli* K-12 across three biological replicates each, yielding nine plates in total. Per-replicate MIC values from all nine plates are summarised in Table 4.

**Table 1. T0 OD₆₀₀ readings for MRSA Rep 1 (19/12/2025).**

| **Row** | **Col 1** | **Col 2** | **Col 3** | **Col 4** | **Col 5** | **Col 6** | **Col 7** | **Col 8** | **Col 9** | **Col 11** | **Col 12** |
| --- | --- | --- | --- | --- | --- | --- | --- | --- | --- | --- | --- |
| A (Nettle) | 1.138 | 0.461 | 0.952 | 0.843 | 0.594 | 0.400 | 0.293 | 0.212 | 0.167 | 0.129 | 0.268 |
| B (Thyme) | 1.979 | 1.623 | 2.019 | 1.691 | 1.074 | 0.683 | 0.474 | 0.326 | 0.185 | 0.198 | 0.164 |
| C (Rosemary) | 2.210 | 2.142 | 2.137 | 2.096 | 1.434 | 0.794 | 0.597 | 0.432 | 0.195 | 0.216 | 0.174 |
| D (Lavender) | 0.250 | 0.361 | 0.356 | 0.361 | 0.343 | 0.345 | 0.356 | 0.335 | 0.225 | 0.184 | 0.157 |
| E (Lemongrass) | 0.350 | 0.363 | 0.373 | 0.362 | 0.362 | 0.376 | 0.365 | 0.338 | 0.294 | 0.179 | 0.155 |
| F (doTERRA) | 0.293 | 0.440 | 0.370 | 0.364 | 0.349 | 0.356 | 0.361 | 0.346 | 0.260 | 0.157 | 0.108 |

*Concentrations across cols 1–8 (mg/mL): 50, 25, 12.5, 6.25, 3.13, 1.56, 0.78, 0.39. Col 10 (additional blank) omitted from this table; values were 0.166, 0.168, 0.172, 0.227, 0.188, 0.231 for rows A–F respectively.*

**Table 2. T24 OD₆₀₀ readings for MRSA Rep 1 (20/12/2025, 24 h post-inoculation).**

| **Row** | **Col 1** | **Col 2** | **Col 3** | **Col 4** | **Col 5** | **Col 6** | **Col 7** | **Col 8** | **Col 9** | **Col 11** | **Col 12** |
| --- | --- | --- | --- | --- | --- | --- | --- | --- | --- | --- | --- |
| A (Nettle) | 0.221 | 0.325 | 0.533 | 0.605 | 0.483 | 0.367 | 0.503 | 0.622 | 0.203 | 0.572 | 0.087 |
| B (Thyme) | 1.605 | 1.441 | 1.995 | 1.592 | 0.959 | 0.680 | 0.428 | 0.770 | 0.183 | 0.341 | 0.089 |
| C (Rosemary) | 1.805 | 2.296 | 2.032 | 2.164 | 1.556 | 0.934 | 0.652 | 0.434 | 0.175 | 0.239 | 0.090 |
| D (Lavender) | 0.260 | 0.287 | 0.291 | 0.275 | 0.266 | 0.236 | 0.738 | 0.586 | 0.164 | 0.125 | 0.088 |
| E (Lemongrass) | 0.580 | 0.642 | 0.291 | 0.255 | 0.235 | 0.211 | 0.260 | 0.168 | 0.141 | 0.102 | 0.088 |
| F (doTERRA) | 0.305 | 0.361 | 0.342 | 0.266 | 0.230 | 0.181 | 0.106 | 0.086 | 0.087 | 0.096 | 0.089 |

**4. Worked Calculation Example**

**Row A (Nettle ethanolic extract) at column 6 (1.56 mg/mL) — the reported MIC for Nettle vs MRSA in Rep 1:**

• T0 (A6) = 0.400; T24 (A6) = 0.367; ΔOD sample = 0.367 − 0.400 = −0.033

• Blank: row A, mean of col 9 and col 12 = mean(0.036, −0.181) = −0.073

• Solvent control (A11): T0 = 0.129; T24 = 0.572; ΔOD = +0.443

• RG = ((−0.033) − (−0.073)) ÷ ((0.443) − (−0.073)) × 100 = 0.040 ÷ 0.516 × 100 = 7.8%

**• 7.8% ≤ 10% threshold → INHIBITED. Column 6 (1.56 mg/mL) is therefore the MIC for this replicate.**

**5. Per-Replicate MIC Summary Across All Nine Plates**

Table 4 below summarises per-replicate MIC values for each preparation × strain combination across all nine biological replicates (three strains × three replicates per strain). Median values reproduce those reported in main manuscript Table 3.

**Table 4. Per-replicate MIC values (mg/mL) across all nine biological replicates.**

| **Preparation** | **Type** | **MRSA R1** | **MRSA R2** | **MRSA R3** | **MRSA Median** | **MSSA R1** | **MSSA R2** | **MSSA R3** | **MSSA Median** | ***E. coli* R1** | ***E. coli* R2** | ***E. coli* R3** | ***E. coli* Median** |
| --- | --- | --- | --- | --- | --- | --- | --- | --- | --- | --- | --- | --- | --- |
| Nettle | EE | 1.56 | 0.39 | 6.25 | 1.56 | 6.25 | 1.56 | 0.39 | 1.56 | 6.25 | 3.13 | 0.39 | 3.13 |
| Thyme | EE | 0.78 | 3.13 | 3.13 | 3.13 | 0.39 | 12.5 | 3.13 | 3.13 | 3.13 | 0.39 | 3.13 | 3.13 |
| Rosemary | EE | 12.5 | 0.39 | 1.56 | 1.56 | 0.39 | 0.39 | 3.13 | 0.39 | 3.13 | 3.13 | 0.39 | 3.13 |
| Lavender | EO | 1.56 | 0.39 | 0.39 | 0.39 | 6.25 | 1.56 | 1.56 | 1.56 | 1.56 | 1.56 | 3.13 | 1.56 |
| Lemongrass | EO | 0.39 | 0.39 | 0.39 | 0.39 | 0.39 | 0.39 | >50* | 0.39 | 0.78 | 50* | 0.78 | 0.78 |
| doTERRA Peace | EO | 0.39 | 0.39 | 6.25 | 0.39 | 0.39 | >50* | 3.13 | 3.13 | 25 | 0.39 | 12.5 | 12.5 |

*EE = ethanolic extract; EO = essential oil. Per-replicate MIC values are the lowest concentration at which relative growth was ≤10% (Subsection 1 formula). Median values reproduce those in main manuscript Table 3. *Anomalous replicates with no inhibition or inhibition only at the highest concentration tested (50 mg/mL): lemongrass MSSA Rep 3 (no inhibition at ≤50 mg/mL), lemongrass E. coli Rep 2 (inhibition only at 50 mg/mL), and doTERRA MSSA Rep 2 (no inhibition at ≤50 mg/mL). These results are interpreted as anomalous per main manuscript Section 3.3; medians are calculated across the three replicates including the anomalous values.*

**Supplementary Material 3**

**Phytochemical Screening Protocols and Notes on GC–MS Interpretability of Ethanolic Extracts**

Detailed protocols for the four qualitative phytochemical screening tests applied to ethanolic extracts of nettle, thyme, and rosemary, and notes on the GC–MS interpretability of these extracts.

**1. Flavonoid Screening (Shinoda Test)**

Reagent: concentrated hydrochloric acid; magnesium turnings. Procedure: 2 mL ethanolic extract was placed in a test tube; a few magnesium turnings were added followed by drop-wise addition of concentrated HCl. Positive indicator: development of pink, crimson, or magenta coloration within 5 minutes. Observed results: nettle and thyme produced yellow-orange and orange-red coloration respectively (recorded as positive); rosemary produced no colour change (recorded as negative). Conducted in triplicate; results consistent across replicates (Harborne, 1998).

**2. Alkaloid Screening (Wagner's Reagent)**

Reagent: 1.27 g iodine and 2 g potassium iodide dissolved in 5 mL distilled water then diluted to 100 mL. Procedure: 2 mL extract was acidified with dilute hydrochloric acid; 2–3 drops of Wagner's reagent were added. Positive indicator: formation of a brown to reddish-brown precipitate within 1 minute. Observed results: all three extracts (nettle, thyme, rosemary) produced brown precipitates (recorded as positive). Conducted in triplicate; results consistent across replicates (Sreevidya and Mehrotra, 2003).

**3. Phenolic Screening (Ferric Chloride Test)**

Reagent: 1% w/v ferric chloride solution in distilled water. Procedure: 2 mL extract was treated with 2–3 drops of 1% ferric chloride solution. Positive indicator: development of blue-black or green coloration. Observed results: all three extracts produced green coloration (recorded as positive). Conducted in triplicate; results consistent across replicates (Dai and Mumper, 2010).

**4. Saponin Screening (Froth Test)**

Procedure: 2 mL extract was diluted with 6 mL distilled water in a test tube and shaken vigorously for 30 seconds. Positive indicator: persistent foam height ≥ 1 cm after 10 minutes. Observed results: no extract produced persistent foam ≥ 1 cm (recorded as negative). Conducted in triplicate; results consistent across replicates (Sofowora, 1993).

**5. Notes on GC–MS Interpretability of Ethanolic Extracts**

Direct-injection GC–MS of nettle, thyme, and rosemary ethanolic extracts produced TIC profiles dominated by early-eluting matrix compounds at retention time 3–6 minutes, without identifiable botanical terpene constituents above background. This is a recognised limitation of GC–MS profiling of complex ethanolic matrices without prior cleanup: the ethanolic solvent and water-soluble polar compounds co-elute with low-molecular-weight matrix constituents and overwhelm the chromatographic signal in the early-eluting region, masking any botanical terpenes that may be present at low relative abundance.

Future characterisation of these preparations should employ either: (a) solid-phase extraction cleanup before GC–MS injection to remove polar matrix constituents and enrich the volatile fraction (Myers et al., 2021), or (b) liquid chromatography–mass spectrometry (LC–MS) using a method capable of resolving polar phenolics and other ethanol-soluble compounds that GC–MS cannot detect. The phytochemical screening results in Subsections 1–4 confirm the presence of phenolic compounds and alkaloids in all three extracts, supporting the case that compositionally informative profiling is possible with appropriate methodology.

**Supplementary Material 4**

**Worked Chemotype Similarity Index Calculation**

Step-by-step worked example demonstrating Chemotype Similarity Index (CSI) calculation for the three essential oil preparations characterised by GC–MS (Sections 2.10 and 2.11 of the main manuscript). The complete Python implementation is available on GitHub at *https://github.com/atulbhat080-crypto/CSI-Framework* and archived on Zenodo with DOI *10.5281/zenodo.20725211*.

**1. Compositional Vector Construction**

For each essential oil profiled by GC–MS, the relative percentage abundance of every compound identified above the NIST 2017 match score threshold of 700 was extracted from the total ion chromatogram and normalised to the total identified compound area, yielding a fractional compositional vector for each preparation. The union of compounds appearing within the top ten constituents of any of the three essential oils was constructed, yielding 23 unique compounds. Compositional vectors for each preparation were aligned across this 23-compound space, with compounds absent from a given preparation assigned a value of zero.

**2. Unified Compound Space (23 Compounds)**

The unified compound space comprises the union of the top ten compounds across lavender, lemongrass, and the doTERRA Peace blend (Table 1 of the main manuscript). Compounds appearing in any preparation above the NIST 2017 HIT 700 threshold are included; compounds not detected in a preparation are assigned a value of zero.

Compounds: α-pinene, camphene, eucalyptol (1,8-cineole), linalool, cis-p-menth-8-en-1-ol, endo-borneol, terpinen-4-ol, (−)-cis-isopiperitenol, citral (neral isomer), linalyl acetate, geraniol, neral, geranial (E-2,6-octadienal), lavandulyl acetate, copaene, cis-α-bisabolene, β-caryophyllene, β-longipinene, germacrene D, caryophyllene oxide, cadinene-type sesquiterpene, aromadendrene-type sesquiterpene, (E)-nerolidol.

**3. Pearson Correlation Calculation**

For each pair of preparations, Pearson product-moment correlation was calculated across the aligned 23-compound vectors:

$$r = (\sum(x\_i - x̄)(y\_i - ȳ))/\sqrt{\sum\left( x_{i}- \bar{x} \right)^{2}\times\sum\left( y_{i}- ȳ \right)^{2}}$$

where xᵢ and yᵢ are the normalised relative abundances of compound i in preparations X and Y respectively, and x̄ and ȳ are the mean abundances across the unified compound space. The calculation yields:

• Lavender vs Lemongrass: r = −0.153 (compositionally unrelated)

• Lavender vs doTERRA Peace: r = +0.716 (strong compositional similarity)

• Lemongrass vs doTERRA Peace: r = −0.256 (compositionally unrelated)

**4. Mean CSI Calculation**

Mean CSI is calculated as the arithmetic mean of the three pairwise Pearson correlations:

*Mean CSI = ((−0.153) + (+0.716) + (−0.256)) / 3 = +0.307 / 3 = +0.102*

A mean CSI of +0.102 indicates that, taken as a panel, the three essential oils are compositionally independent overall, with the strong lavender–doTERRA correspondence balanced by the divergence of lemongrass from both.

**5. Log₂-MIC Difference Calculation**

For each pair of preparations and each bacterial strain, the absolute log₂-MIC difference is calculated as:

$$|\Delta log_{2} \left( MIC \right)| = \left| log_{2} \left( MIC_{a} \right)-log_{2} \left( MIC_{b} \right) \right|$$

Log2 transformation is appropriate because MIC values were generated by two-fold serial dilution; log₂ differences therefore correspond directly to the dilution steps separating preparations.

**Worked example**

Lavender vs doTERRA Peace against *E. coli* K-12: lavender median MIC = 1.56 mg/mL; doTERRA Peace median MIC = 12.5 mg/mL. |Δlog₂MIC| = |log₂(1.56) − log₂(12.5)| = |0.642 − 3.642| = 3.000. This pair therefore shows a 3.0 log₂-step (8-fold) difference in MIC despite a strong positive compositional correlation (+0.716).

**6. Strain-Dependent Chemistry–Activity Analysis**

**Table 1. Pairwise compositional similarity and absolute log₂-MIC differences (reproduces main manuscript Table 8).**

| **Pair** | **Chemical r** | **MRSA \|Δlog₂MIC\|** | **MSSA \|Δlog₂MIC\|** | ***E. coli* \|Δlog₂MIC\|** |
| --- | --- | --- | --- | --- |
| LAV–LEM | −0.153 | 0.00 | 2.00 | 1.00 |
| LAV–dT | +0.716 | 0.00 | 1.00 | 3.00 |
| LEM–dT | −0.256 | 0.00 | 3.00 | 4.00 |

**7. Implementation Notes**

All Pearson correlations and log₂-MIC differences were calculated in Python 3.11 using NumPy 1.26. The complete CSI calculation script with documented inputs, worked example, and reusable function definitions is openly available at *https://github.com/atulbhat080-crypto/CSI-Framework* and archived with a permanent identifier on Zenodo (DOI: *10.5281/zenodo.20725211*) so that the framework may be applied to other compositional datasets. CSI is presented as an exploratory descriptive framework based on n = 3 essential oil preparations; statistical inference on the correlation coefficients is not performed because the sample size precludes adequately powered hypothesis testing.

**Supplementary Material 5**

**Post-Exposure Culturability — CFU/mL Conversions**

Drop-plate enumeration data and log₁₀ CFU mL⁻¹ conversion methodology underpinning Figure 1 in the main manuscript. Drop-plate counts at each timepoint × dilution × replicate were converted to viable counts using the formula below; the most countable dilution was selected per timepoint in accordance with standard drop-plate convention (Herigstad et al., 2001).

**1. Drop-Plate CFU/mL Conversion Formula**

Drop volume = 10 µL (0.01 mL). For each spot count:

$$\frac{CFU}{mL}=\frac{mean valid count}{0.01 mL}\times\frac{1}{dilution factor}$$

where mean valid count = arithmetic mean of three drop replicates per dilution, excluding TMTC (too many to count) and zero values. log₁₀ CFU/mL = log₁₀ of the CFU/mL value.

**2. Worked Example**

**MRSA growth control at 0h (used as reference baseline for all MRSA treatments — see main manuscript Section 3.4):**

• Treatment-well counts at 0h were TMTC at all dilutions tested (10⁻³ to 10⁻⁶), reflecting the high inoculum density (~10⁷ CFU/mL) used in this assay.

• Growth control at 10⁻⁵ dilution gave drop counts: 0, 3, 3 (mean valid = 2 colonies)

• CFU/mL = (2 ÷ 0.01) × 10⁵ = 2.0 × 10⁷

• log₁₀ CFU/mL = log₁₀ (2.0 × 10⁷) = 7.30

**• Reported MRSA 0h baseline = 7.73 log₁₀ CFU mL⁻¹ (mean across all available 0h dilutions including 10⁻⁶).**

**3. Per-Treatment log₁₀ CFU mL⁻¹ Values**

Calculated log₁₀ CFU mL⁻¹ values feeding Figure 1 of the main manuscript, derived from drop-plate counts in the raw MBC dilutions dataset using the 10 µL drop volume conversion in Subsection 1.

**Panel A: MSSA**

**Table 1. log₁₀ CFU mL⁻¹ values across timepoints — MSSA panel.**

| **Treatment** | **0h** | **2h** | **6h** | **24h** |
| --- | --- | --- | --- | --- |
| Growth control | 7.29 | 7.54 | 5.38 | 7.53 |
| Vancomycin | 6.99 | 7.70 | 5.37 | 7.18 |
| Lavender 1× MIC | 6.11 | 6.08 | 6.00 | 7.19 |
| Lavender 2× MIC | 7.04 | 5.95 | 6.53 | 7.54 |
| Rosemary 2× MIC | 7.88 | 7.32 | 6.60 | TMTC* |

**TMTC (Too many to count) at all dilutions tested at 24h — excluded from main manuscript Figure 1.*

**Panel B: MRSA**

**Table 2. log₁₀ CFU mL⁻¹ values across timepoints — MRSA panel.**

| **Treatment** | **0h** | **2h** | **6h** | **24h** |
| --- | --- | --- | --- | --- |
| Growth control | 7.73 | 6.78 | 6.74 | 7.12 |
| Vancomycin | 7.73 | 8.95** | 4.90 | 6.60 |
| Lavender 1× MIC | 7.73 | 7.26 | 5.19 | 7.30 |
| Lavender 2× MIC | 7.73 | 5.30 | 5.30 | 7.30 |
| Thyme 2× MIC | 7.73 | 7.18 | 6.32 | 7.50 |

*Treatment wells were TMTC at 0h; the GC 0h value (7.73) is used as the reference baseline for all treatments. **Vancomycin 2h (8.95) excluded as anomalous artefact inconsistent with adjacent timepoints.*

**Panel C: *E. coli* K-12**

**Table 3. log₁₀ CFU mL⁻¹ values across timepoints — *E. coli* K-12 panel.**

| **Treatment** | **0h** | **2h** | **6h** | **24h** |
| --- | --- | --- | --- | --- |
| Growth control | 7.19 | 6.15 | 8.04 | TMTC* |
| Ciprofloxacin | <2.0 | 6.18 | 4.78 | <2.0 |
| Lavender 1× MIC | 7.02 | 6.29 | 5.93 | 7.40 |
| Lavender 2× MIC | 6.27 | 5.54 | TMTC* | 7.48 |
| Thyme 2× MIC | 7.20 | 6.95 | 8.26 | 7.28 |

**TMTC ( too many to count) at all dilutions tested. Ciprofloxacin counts below the drop-plate detection limit of approximately 2 log₁₀ CFU mL⁻¹ at 0h and 24h.*

**4. Notes on Interpretation**

Reductions are reported relative to the growth control 0h value of each panel (MSSA 7.29, MRSA 7.73, *E. coli* 7.19) for consistency across treatments. The bactericidal threshold of ≥3 log₁₀ CFU mL⁻¹ reduction (Wiegand et al., 2008) was not reached by any preparation at any timepoint. The largest reduction observed was lavender 2× MIC vs MRSA at 2h (2.43-log reduction; baseline 7.73 → 5.30); all reductions recovered toward baseline by T24, consistent with bacteriostatic rather than bactericidal activity.

**5. Inoculum Density Verification**

OD₆₀₀ readings of the standardised 0.5 McFarland suspensions used for this assay ranged from 0.08 to 0.10 against a BHI blank, consistent with the conventional 0.5 McFarland turbidity standard. This corresponds to approximately 10⁸ CFU/mL pre-dilution; the 1:10 dilution into the assay therefore yielded a starting inoculum of approximately 10⁷ CFU/mL — approximately 20-fold higher than the inoculum used in the MIC assay (main manuscript Section 2.6). This higher inoculum density is consistent with time-kill assay convention and explains the elevated GC 0h baseline values across all three panels (7.19–7.73 log₁₀ CFU mL⁻¹).

**Supplementary Material 6**

**Antibiotic MIC Determination**

Vancomycin and ciprofloxacin MIC values used as denominator references for the checkerboard FICI assay (Section 3.5 of the main manuscript) were determined by broth microdilution in BHI broth on a single 96-well plate (15–16 April 2026). The data presented below document the relative growth (RG) calculations underpinning the FICI denominators reported in the main text. This experiment was conducted once (n = 1) with duplicate rows per condition; this single-experiment status is acknowledged as a methodological limitation and is the basis for the discussion in Section 4.5 of the main manuscript regarding inoculum-mediated MIC shift.

**1. Relative Growth Formula**

Relative growth (RG) was calculated for every well as:

$$RG \left( \% \right)=\frac{\Delta OD_{sample}- \Delta OD_{blank}}{\Delta OD_{growth control}- \Delta OD_{blank}}\times100$$

where ΔOD = OD₆₀₀ at 24h minus OD₆₀₀ at 0h for each well; blank = column 12 (BHI broth, no bacteria, no antibiotic); growth control = column 9 (bacteria in BHI broth, no antibiotic). The inhibition threshold was set at RG ≤ 10%.

**2. Plate Layout**

The single 96-well plate was loaded as follows:

| **Rows** | **Antibiotic** | **Strain** | **Notes** |
| --- | --- | --- | --- |
| A and B | Vancomycin | MRSA | Duplicate rows; same dilution series |
| C and D | Vancomycin | MSSA | Duplicate rows; same dilution series |
| E and F | Ciprofloxacin | *E. coli* K-12 | Row F GC failed — excluded from MIC determination |
| Col 9 | Growth control | All | Bacteria in BHI, no antibiotic |
| Col 12 | Blank | — | BHI broth only, no bacteria, no antibiotic |

Antibiotic concentrations across columns 1–8 (two-fold serial dilution from column 1):

| **Antibiotic** | **Col 1** | **Col 2** | **Col 3** | **Col 4** | **Col 5** | **Col 6** | **Col 7** | **Col 8** |
| --- | --- | --- | --- | --- | --- | --- | --- | --- |
| Vancomycin (µg/mL) | 30 | 15 | 7.5 | 3.75 | 1.88 | 0.94 | 0.47 | 0 |
| Ciprofloxacin (µg/mL) | 0.5 | 0.25 | 0.125 | 0.063 | 0.031 | 0.016 | 0.008 | 0 |

**3. Raw Absorbance Data**

**Table 1. 0h OD₆₀₀ readings (15/04/2026, 16:27).**

| **Row** | **Col 1** | **Col 2** | **Col 3** | **Col 4** | **Col 5** | **Col 6** | **Col 7** | **Col 8** | **Col 9** | **Col 12** |
| --- | --- | --- | --- | --- | --- | --- | --- | --- | --- | --- |
| A | 0.090 | 0.086 | 0.094 | 0.092 | 0.093 | 0.090 | 0.095 | 0.095 | 0.093 | 0.099 |
| B | 0.094 | 0.089 | 0.092 | 0.091 | 0.093 | 0.090 | 0.090 | 0.094 | 0.091 | 0.098 |
| C | 0.088 | 0.092 | 0.091 | 0.093 | 0.091 | 0.090 | 0.091 | 0.090 | 0.092 | 0.098 |
| D | 0.093 | 0.089 | 0.090 | 0.090 | 0.091 | 0.094 | 0.096 | 0.090 | 0.091 | 0.100 |
| E | 0.090 | 0.091 | 0.115 | 0.091 | 0.089 | 0.089 | 0.091 | 0.114 | 0.093 | 0.099 |
| F | 0.092 | 0.091 | 0.088 | 0.094 | 0.090 | 0.091 | 0.092 | 0.105 | 0.092 | 0.098 |

**Table 2. 24h OD₆₀₀ readings (16/04/2026, 16:43).**

| **Row** | **Col 1** | **Col 2** | **Col 3** | **Col 4** | **Col 5** | **Col 6** | **Col 7** | **Col 8** | **Col 9** | **Col 12** |
| --- | --- | --- | --- | --- | --- | --- | --- | --- | --- | --- |
| A | 0.317 | 0.792 | 0.706 | 0.729 | 0.740 | 0.739 | 0.943 | 0.907 | 0.787 | 0.273 |
| B | 0.293 | 0.692 | 0.721 | 0.847 | 0.985 | 0.757 | 0.790 | 0.818 | 0.892 | 0.314 |
| C | 0.240 | 0.688 | 0.827 | 0.906 | 0.759 | 0.914 | 0.888 | 0.809 | 0.924 | 0.308 |
| D | 0.130 | 0.572 | 0.692 | 0.749 | 0.848 | 0.913 | 0.826 | 0.660 | 0.882 | 0.322 |
| E | 0.086 | 0.333 | 0.554 | 0.755 | 0.609 | 0.502 | 0.605 | 1.235 | 0.887 | 0.284 |
| F | 0.091 | 0.086 | 0.232 | 0.516 | 0.553 | 0.419 | 0.507 | 1.237 | 0.279 | 0.099 |

**4. Worked Calculation Example**

**Row B, Column 1 (Vancomycin 30 µg/mL vs MRSA):**

• ΔOD sample (B1) = 0.293 − 0.094 = 0.1981

• ΔOD blank (B12) = 0.314 − 0.098 = 0.2163

• ΔOD growth control (B9) = 0.892 − 0.091 = 0.8009

• RG = (0.1981 − 0.2163) ÷ (0.8009 − 0.2163) × 100 = −3.1%

**• −3.1% ≤ 10% threshold → INHIBITED. Column 1 (Vancomycin 30 µg/mL) inhibits MRSA in row B.**

**5. Full Relative Growth Tables**

**Table 3. Vancomycin RG (%) vs MRSA (rows A–B) and MSSA (rows C–D).**

*Inhibited cells (RG ≤ 10%) are flagged with green shading. RG values calculated per well using row-specific blank correction.*

| **Row** | **30 µg/mL** | **15** | **7.5** | **3.75** | **1.88** | **0.94** | **0.47** | **0 (no AB)** | **GC ΔOD** | **BLK ΔOD** |
| --- | --- | --- | --- | --- | --- | --- | --- | --- | --- | --- |
| A (Van vs MRSA) | 10.1% | 102.3% | 84.2% | 89.2% | 90.9% | 91.5% | 129.8% | 122.8% | 0.694 | 0.174 |
| B (Van vs MRSA) | −3.1% | 66.3% | 70.5% | 92.3% | 115.6% | 77.1% | 82.7% | 86.8% | 0.801 | 0.216 |
| C (Van vs MSSA) | −9.4% | 62.0% | 84.5% | 96.9% | 73.6% | 98.7% | 94.3% | 81.7% | 0.832 | 0.210 |
| D (Van vs MSSA) | −32.6% | 46.0% | 66.8% | 76.7% | 94.1% | 104.8% | 89.2% | 61.1% | 0.791 | 0.222 |

*Row A col 1 (10.1%) is flagged as borderline at the threshold; rows B, C, D col 1 (≤0%) flagged as clearly inhibited.*

**Table 4. Ciprofloxacin RG (%) vs *E. coli* K-12 (rows E–F).**

*Row F is shown for completeness but excluded from MIC determination because the growth control failed (GC ΔOD = 0.187 vs row E GC ΔOD = 0.794). All row F RG values asterisked are mathematically unreliable owing to the failed growth control.*

| **Row** | **0.5 µg/mL** | **0.25** | **0.125** | **0.063** | **0.031** | **0.016** | **0.008** | **0 (no AB)** | **GC ΔOD** | **BLK ΔOD** |
| --- | --- | --- | --- | --- | --- | --- | --- | --- | --- | --- |
| E (Cipro vs *E. coli*) | −31.2% | 9.4% | 41.6% | 78.7% | 55.0% | 37.4% | 54.2% | 153.9% | 0.794 | 0.185 |
| F (Cipro vs *E. coli*, excluded) | −0.6%* | −2.8%* | 76.6%* | 225.6%* | 248.0%* | 175.1%* | 221.8%* | 605.2%* | 0.187 | 0.000 |

**6. MIC Summary**

**Table 5. Summary of antibiotic MIC values determined from this single experiment.**

| **Antibiotic** | **Strain** | **MIC** | **Notes** |
| --- | --- | --- | --- |
| Vancomycin | MRSA | 30 µg/mL | Inhibited at col 1 only. True MIC 15–30 µg/mL — upper boundary of tested range. |
| Vancomycin | MSSA | 30 µg/mL | Inhibited at col 1 only. Concordant across rows C and D. |
| Ciprofloxacin | *E. coli* K-12 | 0.25 µg/mL | Based on row E only. Row F excluded — failed GC. |

All MIC determinations were performed in BHI broth, consistent with all other antimicrobial assays in this study. Values may differ from CAMHB reference ranges; this is consistent with documented medium-dependent variability in broth microdilution susceptibility testing (Wiegand et al., 2008). The possibility of an inoculum-mediated upward shift in vancomycin MIC is discussed in Section 4.5 of the main manuscript.

**Supplementary Material 7**

**Checkerboard FICI Synergy Assay**

Per-plate validity assessment and FICI calculation methodology for the four two-compound combinations described in Section 3.5 of the main manuscript. Twelve 96-well plates were assessed in total: four combinations × three biological replicates each. All plates were prepared and inoculated on the same day and incubated at 37 °C for 24 hours.

**1. Combinations Tested**

**Table 1. The four checkerboard combinations and their parameters.**

| **Combo** | **Plant Preparation** | **Antibiotic** | **Strain** | **Highest extract** |
| --- | --- | --- | --- | --- |
| 1 | Lavender EO | Vancomycin | MRSA | 0.78 mg/mL (2× MIC) |
| 2 | Lemongrass EO | Vancomycin | MRSA | 0.78 mg/mL (2× MIC) |
| 3 | Rosemary EE | Vancomycin | MSSA | 0.78 mg/mL (2× MIC) |
| 4 | Lavender EO | Ciprofloxacin | *E. coli* K-12 | 1.56 mg/mL (1× MIC)* |

**Combination 4 capped at 1× MIC due to the 1% (v/v) DMSO solubility threshold; preparation at 2× MIC would have required 1.42% DMSO in the final well.*

**2. Plate Layout**

Each 96-well plate was configured identically:

| **Region** | **Content** |
| --- | --- |
| Cols 1–7, rows A–G | Combination matrix: antibiotic × extract two-fold serial dilutions |
| Col 8, rows A–G | Extract-alone reference (no antibiotic) |
| Row H, cols 1–7 | Antibiotic-alone reference (no extract) |
| Col 9 | Growth control (bacteria in BHI, no treatment) |
| Col 10 | Antibiotic control (2× MIC + bacteria) |
| Col 11 | Extract control (1× MIC + bacteria) |
| Col 12 | Blank (BHI broth only, no bacteria, no treatment) |

**2a. Antibiotic Working-Stock Preparation**

Antibiotic working stocks were prepared above the target final concentration to account for in-well dilution. Each final matrix well contained 50 µL antibiotic + 50 µL extract + 10 µL inoculum (110 µL total), giving a dilution factor of 110/50 = 2.2×. The vancomycin working stock was therefore 132 µg/mL (column-1 final = 60 µg/mL = 2× the 30 µg/mL standalone MIC) and the ciprofloxacin working stock was 1.10 µg/mL (column-1 final = 0.50 µg/mL = 2× the 0.25 µg/mL MIC). Columns 1–7 represent two-fold serial dilutions of these final concentrations (vancomycin 60 → 0.94 µg/mL; ciprofloxacin 0.50 → 0.008 µg/mL).

**3. FICI Calculation Formula**

The Fractional Inhibitory Concentration Index was calculated as:

$$FICI = MIC_{\text{extract, combination}}/MIC_{\text{extract, alone}} + MIC_{\text{antibiotic, combination}}/MIC_{\text{antibiotic, alone}}$$

Classification (Odds, 2003): ≤ 0.5 synergy; 0.5–1.0 additive; 1.0–4.0 indifference; > 4.0 antagonism. Relative growth (RG) was calculated for every well as in SM 6 Subsection 1, with column 9 as the growth control reference and column 12 as the row blank. Wells were classified as inhibited at RG ≤ 10%. Rows where the column 12 blank ΔOD > 0.2 were flagged as contaminated and excluded.

**4. Plate Validity Audit**

**Table 2. Growth control validity across all 12 plates.**

*ΔOD growth control values across rows A–H per replicate. Mean ΔOD GC across all valid rows was 1.08 (range 0.91–1.26), confirming bacterial growth on every plate.*

| **Combination** | **Rep 1 GC range** | **Rep 2 GC range** | **Rep 3 GC range** |
| --- | --- | --- | --- |
| C1: Lavender + Vanco vs MRSA | 1.08–1.26 | 1.07–1.20 | 1.06–1.20 |
| C2: Lemongrass + Vanco vs MRSA | 1.07–1.15 | 1.05–1.13 | 0.97–1.09 |
| C3: Rosemary + Vanco vs MSSA | 1.06–1.16 | 1.03–1.15 | 0.97–1.08 |
| C4: Lavender + Cipro vs *E. coli* | 1.07–1.13 | 0.93–1.15 | 0.91–1.05 |

All 12 plates produced GC ΔOD values ≥ 0.91, well above the conventional viability threshold of 0.30 (Wiegand et al., 2008).

**5. Reference Control Inhibition Assessment**

**Table 3. Antibiotic-alone (row H) and extract-alone (col 8) reference control RG values at the highest concentration tested.**

*RG ≥ 10% indicates no inhibition at that concentration.*

| **Combo** | **Antibiotic at 2× MIC** | **Extract at highest** | **FICI calculable?** |
| --- | --- | --- | --- |
| C1 | RG 42–96% (no inhibition at any rep) | RG 93–98% at 0.78 mg/mL | No |
| C2 | RG 65–104% (no inhibition at any rep) | RG 95–104% at 0.78 mg/mL | No |
| C3 | RG 82–99% (no inhibition at any rep) | RG 82–99% at 0.78 mg/mL | No |
| C4 | Cipro inhibition at 0.5 µg/mL: RG 3–15% (rep 1, 3); 12–29% (rep 2) | RG 82–105% at 1.56 mg/mL | No — extract inhibition boundary not reached |

In combinations 1–3, vancomycin failed to demonstrate concentration-dependent inhibition at 2× MIC under checkerboard conditions despite the same vancomycin stock producing complete inhibition in standalone MIC plates (SM 6). This pattern is consistent with an inoculum-mediated upward shift in vancomycin MIC and is discussed in Section 4.5 of the main manuscript.

**6. Worked FICI Calculation (Illustrative)**

Although no FICI value in this study was formally calculable, the calculation procedure is illustrated below for transparency. If lavender at 0.39 mg/mL combined with ciprofloxacin at 0.0625 µg/mL had produced inhibition (RG ≤ 10%), the FICI would be:

• MIC ext,combination = 0.39 mg/mL (a sub-MIC matrix concentration, ¼ of the 1.56 mg/mL E. coli MIC)

• MIC ext,alone (main manuscript Section 3.3) = 1.56 mg/mL

• MIC ab,combination = 0.0625 µg/mL

• MIC ab,alone (SM 6) = 0.25 µg/mL

**• FICI = (0.39 ÷ 1.56) + (0.0625 ÷ 0.25) = 0.25 + 0.25 = 0.50 → SYNERGY (boundary)**

No combination-matrix well in any of the 12 plates produced inhibition matching this pattern; all four combinations were therefore classified as 'indifference'.

**7. Contamination Exclusions**

**Table 4. Per-replicate contamination exclusions.**

*Individual well contamination was identified by elevated col 12 blank ΔOD (> 0.2) and affected rows were excluded from RG calculations.*

| **Plate** | **Replicate** | **Excluded rows** | **Reason** |
| --- | --- | --- | --- |
| C1 | Rep 1 | Row A | Blank ΔOD = 0.974 (growth in well A12 with no bacteria) |
| C2 | Rep 1 | Row C | Blank ΔOD = 0.311 |
| C2 | Rep 2 | Row B | Blank ΔOD = 0.947 |
| C2 | Rep 3 | Row D | Blank ΔOD = 0.451 |
| C4 | Rep 1 | Row C | Blank ΔOD = 1.048 |
| C4 | Rep 2 | Rows B, D, F, G | Blank ΔOD = 0.993, 0.969, 0.264, 0.267 |
| C4 | Rep 3 | Rows B, E | Blank ΔOD = 0.828, 0.845 |

All exclusions were applied consistently and are flagged in the corresponding figures (main manuscript Figures 2-5).
